# Uncalled4 improves nanopore DNA and RNA modification detection via fast and accurate signal alignment

**DOI:** 10.1101/2024.03.05.583511

**Authors:** Sam Kovaka, Paul W. Hook, Katharine M. Jenike, Vikram Shivakumar, Luke B. Morina, Roham Razaghi, Winston Timp, Michael C. Schatz

## Abstract

Nanopore signal analysis enables detection of nucleotide modifications from native DNA and RNA sequencing, providing both accurate genetic/transcriptomic and epigenetic information without additional library preparation. Presently, only a limited set of modifications can be directly basecalled (e.g. 5-methylcytosine), while most others require exploratory methods that often begin with alignment of nanopore signal to a nucleotide reference. We present Uncalled4, a toolkit for nanopore signal alignment, analysis, and visualization. Uncalled4 features an efficient banded signal alignment algorithm, BAM signal alignment file format, statistics for comparing signal alignment methods, and a reproducible *de novo* training method for k-mer-based pore models, revealing potential errors in ONT’s state-of-the-art DNA model. We apply Uncalled4 to RNA 6-methyladenine (m6A) detection in seven human cell lines, identifying 26% more modifications than Nanopolish using m6Anet, including in several genes where m6A has known implications in cancer. Uncalled4 is available open-source at github.com/skovaka/uncalled4.

## Introduction

Long-read single-molecule sequencers from Oxford Nanopore Technologies (ONT) and Pacific Biosciences (PacBio) have increasing utility in generating complete genomes and transcriptomes by improving resolution of complex DNA and RNA sequences [1–3]. These sequencers can also detect nucleotide modifications without any specialized library preparation, enabling genome-wide epigenetic profiling including within highly repetitive regions that could not be accurately aligned to with short-reads [4]. Nanopore sequencing is unique in not relying on sequencing-by-synthesis, instead measuring electric current that varies over time as nucleotides pass through a pore. While many analyses only use the basecalled sequence, inclusion of the electric current can improve fidelity in several applications, including error correction [5,6], real-time targeted sequencing [7,8], and nucleotide modification detection [9]. Furthermore, ONT is currently the only commercially available platform for directly sequencing RNA without generation of complementary DNA (cDNA), enabling detection of epitranscriptomic modifications. Over 150 known RNA modifications are known to exist, although only a few have been detected thus far, with varying accuracy [10].

Early nanopore sequencers exhibited a high error rate, which could be improved via signal-based polishing [5] or advanced basecalling algorithms. However, a combination of improvements to sequencing chemistry and computational methods have decreased the average ONT DNA sequencing error rate to near 1%, making signal-based polishing largely unnecessary for DNA. This was achieved, in part, by a recent major DNA chemistry update to the r10.4.1 pore, which features two “reader heads” rather than the one present in the previous standard, r9.4.1 (**Fig. 1a**). Direct RNA accuracy has lagged behind, where signal-based polishing can still improve splice site identification [11]. On the software side, modern basecallers use neural networks trained on known sequences to translate the electrical signal into nucleotide reads, with the network architecture and training data being major factors in their accuracy [12]. However, ONT basecallers do not provide an accurate mapping between individual bases and the signal segments which represent them, instead requiring signal alignment to obtain such a mapping.

The latest ONT basecallers can directly detect 5-methylcytosine (5mC) and 5-hydroxymethylcytosine (5hmC) from individual DNA reads [13]. 5mC is the most common DNA modification in humans, with wide ranging effects in cellular development usually via suppression of transcription, with aberrant DNA methylation a frequent hallmark of several types of cancer [14,15]. Progress has also been reported from ONT on direct RNA basecalling for 6-methyladenine (m6A), the most common RNA modification in humans, using the early access RNA004 sequencing chemistry, currently limited to DRACH motifs (D = A, G or T; R = A or G; H=A, C or T) where over 60% of m6A sites occur in humans [16]. M6a is one of several known epitranscriptomic modifications, with diverse effects including transcript decay [17,18], transcript stabilization [19,20], and increased translational efficiency [21]. Beyond 5mC, 5hmC, and m6A in limited contexts, detection of other modifications requires specialized computational methods run after basecalling. Some methods infer modifications from basecaller errors, but these are highly derived results that are sensitive to changes in basecalling models [10], while the most advanced methods analyze the raw signal directly, usually in combination with basecalled reads and beginning with alignment between the signal and a nucleotide reference.

Nanopore signal alignment (also known as event alignment [22,23], segmentation [9], and resquiggling [24]) is analogous to standard nucleotide alignment in variant calling, where the read aligner is often separate from the variant caller. This is also the case for many RNA modification callers which rely on Nanopolish [22] or Tombo [24], the two most commonly used signal aligners [10]. Signal alignment begins by translating the reference sequence into electrical current using a k-mer pore model, which maps each k-mer to the current expected when that combination of bases are in the pore. Processed read signal (e.g. normalized) is then aligned to the expected reference current using a dynamic programming algorithm, such as Dynamic Time Warping (DTW, used by Tombo) or Hidden Markov Models (HMMs, used by Nanopolish). The different methods generally produce similar, although slightly different alignments, with low-complexity sequences or noisy signal often resulting in large-scale disagreements. Comparisons between the methods are hindered by the lack of a ground truth or a shared file format, causing most modification detectors to only support one aligner. More substantially, Nanopolish and Tombo have also not been updated for certain changes to ONT software and chemistry: neither supports the new r10.4.1 DNA chemistry or POD5 signal format, and Tombo relies on deprecated single FAST5 files.

Here we present Uncalled4, a software toolkit for nanopore signal alignment, analysis, and visualization (**Fig. 1**). Uncalled4 features command line tools and interactive visualizations for signal-to-reference alignments, methods for training new pore models, comparisons between alignment methods, and modification level statistics. Uncalled4 uses a banded alignment algorithm guided by basecaller metadata, making it several times faster than Nanopolish or Tombo, and outputs an efficient and indexable BAM file which is directly convertible to widely supported and human-readable file formats. We use Uncalled4 to train a DNA pore model for the r10.4.1 pore, and apply this model to CpG 5mC detection. We also show Uncalled4 outperforms Nanopolish and Tombo in RNA modification detection using several different detection methods. We apply Uncalled4 and m6Anet to seven normal and cancer human cell lines and find 26% more sites supported by the m6A-Atlas than Nanopolish with equivalent precision, highlighting increased sensitivity in several genes with known m6A-related functions. Uncalled4 is implemented in C++ and Python, and is available open source at github.com/skovaka/uncalled4

**Figure 1.**
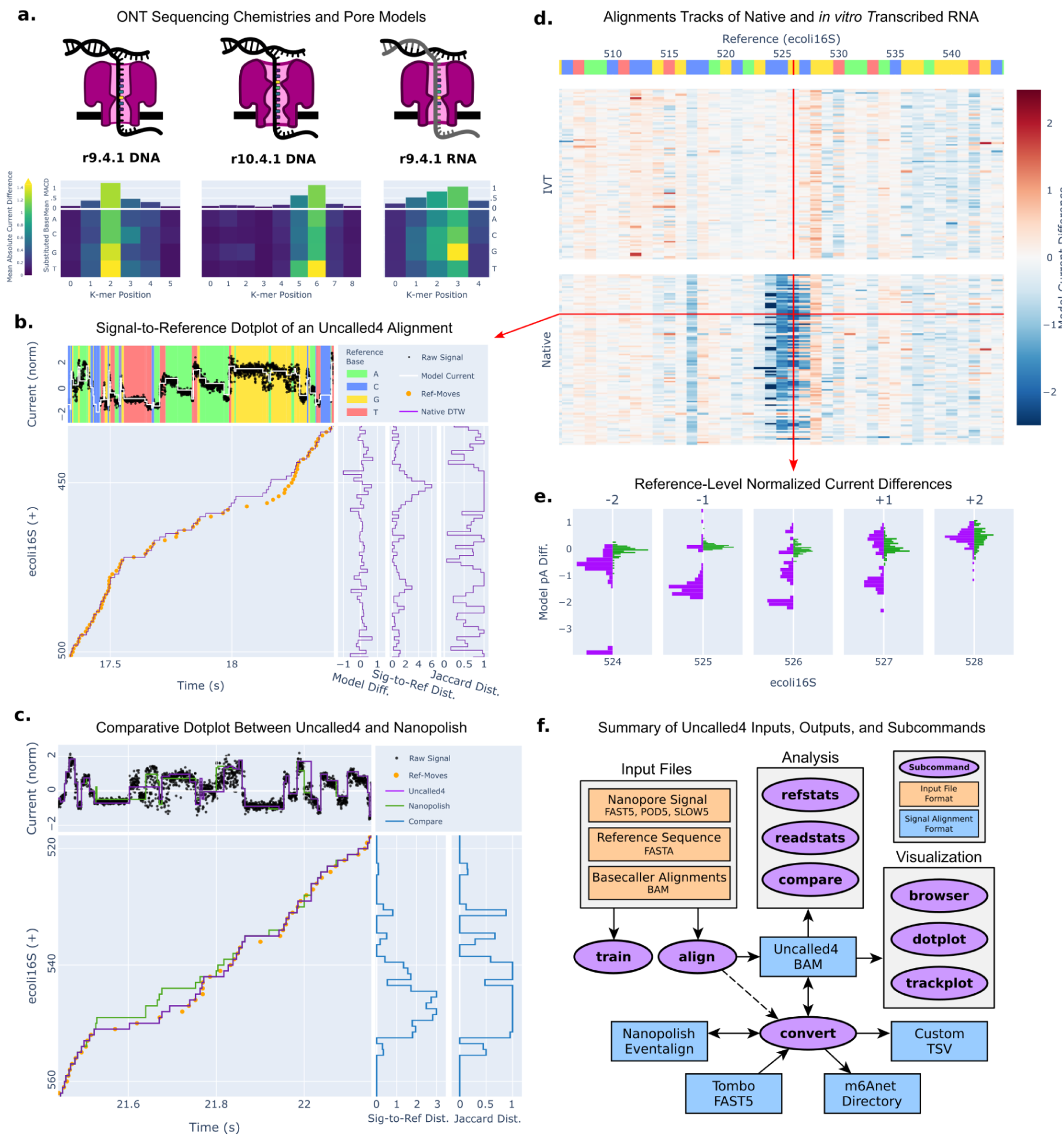
Pore model and alignment methods overview. **(a)** Schematics of Nanopore sequencing chemistries and their pore k-mer substitution profiles. Heatmaps show the mean normalized current difference observed by substituting each base (y-axis) at each k-mer position (x-axis) averaged over all k-mers in the model. **(b)** A signal-to-reference dotplot of an *Escherichia coli* 16S ribosomal RNA (rRNA) read sequenced using ONT r9.4 direct RNA sequencing. Top panel shows the raw samples (black) plotted over the reference base it was aligned to, with the expected pore model current in white. Main panel shows the Uncalled4 read alignment (purple line) over the projected basecaller metadata alignment (orange dots). Side panels show per-reference coordinate summary statistics for the alignment. **(c)** A comparative signal-to-reference dotplot and distance metrics between the alignments. **(d)** A trackplot displaying heatmaps of many native (top) and *in vitro* transcribed (IVT, bottom) *E. coli* 16S rRNA reads aligned by Uncalled4, colored by the difference between the observed and expected normalized current level. Top bar is colored by reference base, and an O6-methylguanine site is known to occur at position 526. **(e)** A refplot summarizing the distributions of differences between observed and expected normalized current levels for native (purple) and IVT (green) reads. **(f)** Schematic of Uncalled4 inputs, outputs, and subcommands (see **Methods**).

## Results

### Alignment efficiency, accuracy, and visualization

Analogous to Nanopore read mapping, Nanopore signal alignment produces a mapping between stretches of nanopore electrical current and reference nucleotides. Conventional nucleotide alignments of basecalled reads (i.e. from minimap2 [25]) determine the coordinates of the reference sequence, which is encoded into expected current using a k-mer pore model. Each pore model is specific to a particular sequencing chemistry, defined by molecule type (RNA or DNA), pore version (e.g. r9.4.1, r10.4.1), sequencing speed (e.g. 400bps), and the output nucleotides, including possibly one or more modifications (e.g. 5mCpG). The number of nucleotides affecting the current level varies by pore, with r10.4’s double reader head necessitating longer k-mers than r9.4. This can be quantified by computing its “substitution profile”: the average normalized current change observed by substituting each base at each k-mer position (**Fig. 1a**). We call the k-mer position with the most influence on current level the “central base”, and observe generally less influence at positions further from the central base, with the exception of r10.4.1 where a secondary reader head generates a smaller, but consistent secondary effect near the beginning of the k-mer (**Fig. 1a**).

**Table 1.** Alignment time, disk space usage, and distance from projected basecaller alignments for Uncalled4 and comparable Nanopore signal aligners. Tombo stores signal alignments alongside raw signal data in single FAST5 files, so space usage is reported as “additional space”/”total FAST5 size”.

|  |  | File size<br>(MB) | Time<br>per 4k<br>reads | Median<br>moves<br>dist | Mean<br>moves<br>dist | Median<br>Jaccard | Mean<br>Jaccard | Mean<br>cov frac |
| --- | --- | --- | --- | --- | --- | --- | --- | --- |
| r9.4 RNA<br>(HEK293T) | Uncalled4 | 31.97 | 113.9 | 0.571 | 0.6701 | 0.7222 | 0.6711 | 98.73% |
|  | Nanopolish | 737.50 | 194.9 | 0.533 | 0.8715 | 0.7143 | 0.6670 | 90.71% |
|  | f5c | 737.39 | 144.7 | 0.533 | 0.8702 | 0.7143 | 0.6670 | 90.71% |
|  | Tombo | 95.07 /<br>254.95 | 772.8 | 0.688 | 14.4741 | 0.8182 | 0.7287 | 98.31% |
| r9.4 DNA | Uncalled4 | 131.04 | 350.6 | 0.375 | 0.8351 | 0.6000 | 0.5958 | 96.64% |
|  | Nanopolish | 3212.52 | 653.8 | 0.375 | 1.0436 | 0.5833 | 0.5864 | 85.97% |
|  | f5c | 3212.52 | 530.0 | 0.667 | 1.044 | 0.5833 | 0.5864 | 85.97% |
|  | Tombo | 389.45 /<br>640.26 | 1015.0 | 2.333 | 55.3012 | 1.0000 | 0.9621 | 76.34% |
| r10.4.1<br>DNA | Uncalled4 | 132.25 | 244.7 | 0.615 | 0.996 | 0.7368 | 0.7116 | 97.38% |
|  | f5c | 3706.41 | 574.4 | 0.654 | 4.734 | 0.7500 | 0.7227 | 89.46% |

Uncalled4 uses basecaller-guided dynamic time warping (bcDTW) to rapidly and accurately align nanopore electrical signal, guided by basecaller metadata (“moves”) in addition to conventional read alignments (**Fig. 1b, Supplemental Figure 1**). By limiting the alignment search space to a narrow range defined by projecting *moves* into reference coordinates (“ref-moves”), Uncalled is substantially faster than Tombo (2.9-6.8x), Nanopolish (1.7-1.9x), and f5c (1.3-2.7x), which is a GPU implementation of Nanopolish that outputs nearly identical alignments [23] (**Table 1**). The computational performance of Uncalled4 varies by molecule and pore type, with direct RNA being fastest per-read due to short read lengths. Uncalled4 encodes signal alignments in BAM files, a compressed format several times smaller than alternatives, e.g. 20 times smaller than Nanopolish’s eventalign format (**Table 1**). Uncalled4 also supports read signal input from FAST5, SLOW5/BLOW5 [26], and POD5 files, the latter of which is the new ONT standard and not supported by Nanopolish or Tombo.

We assess the similarity between each alignment method using two different measurements of pairwise “distance” between alignment paths: *signal Jaccard distance*, defined as the fraction of raw samples that both methods aligned to the same position, and *signal-to-reference distance*, defined as the average nucleotide distance between reference coordinates over each raw sample for each method (**Fig. 1b**). We use these metrics to compare Uncalled4, Nanopolish/f5c, and Tombo alignments of DNA from *Drosophila melanogaster* (r9.4.1 and r10.4.1) and RNA from the human HEK293T cell line (RNA002) (**Supplemental Table 1**). Uncalled4 is most similar to Nanopolish for r9.4.1 DNA and RNA with a median distance of zero in both metrics, although higher mean distances indicate outliers from large-scale discordance in certain reads (**Supplemental Figure 2**). f5c is the only other aligner to support r10.4.1 DNA alignment, which also has zero median distance to Uncalled4, but also shows outlier signals. Tombo is more divergent, with fewer than half of all reference coordinates matching Uncalled4 or Nanopolish exactly. It is notable that the distances are generally greater for DNA than RNA, despite the lower error rate in DNA sequencing. This is largely due to intergenic repeats not present in RNA, which are less likely to disrupt Uncalled4 alignments due to the basecaller guided DTW algorithm (**Supplemental Figure 2a**). Large-scale alignment disagreements are also often found near alignment endpoints, near deletions or insertions, or when the molecule speed changes (**Supplemental Figure 2b-d**).

Distance can also be measured between each alignment method and the *ref-moves* used to guide Uncalled4 bcDTW alignment, generally yielding higher distances due to the low-resolution *moves* (**Fig.1c, Supplemental Table 1**). Uncalled4 consistently has the lowest mean signal-to-reference distance to *ref-moves*. Nanopolish has slightly lower average Jaccard distance in r9.4.1 DNA and RNA, with less than 50% overlap on average for all aligners and less than 2% difference between Uncalled4 and Nanopolish. This can likely be explained by the widespread masking performed by Nanopolish, with 9.3% of positions masked by Nanopolish compared to only 1.3% by Uncalled4. This masking is intended to eliminate noisy signal from Nanopolish alignments, but it also reduces the sensitivity of alignments, especially surrounding modifications which increase the apparent noise in the signal. This is most apparent at low sequencing coverage, where Nanopolish has reduced recall (see below).

**Figure 2.**
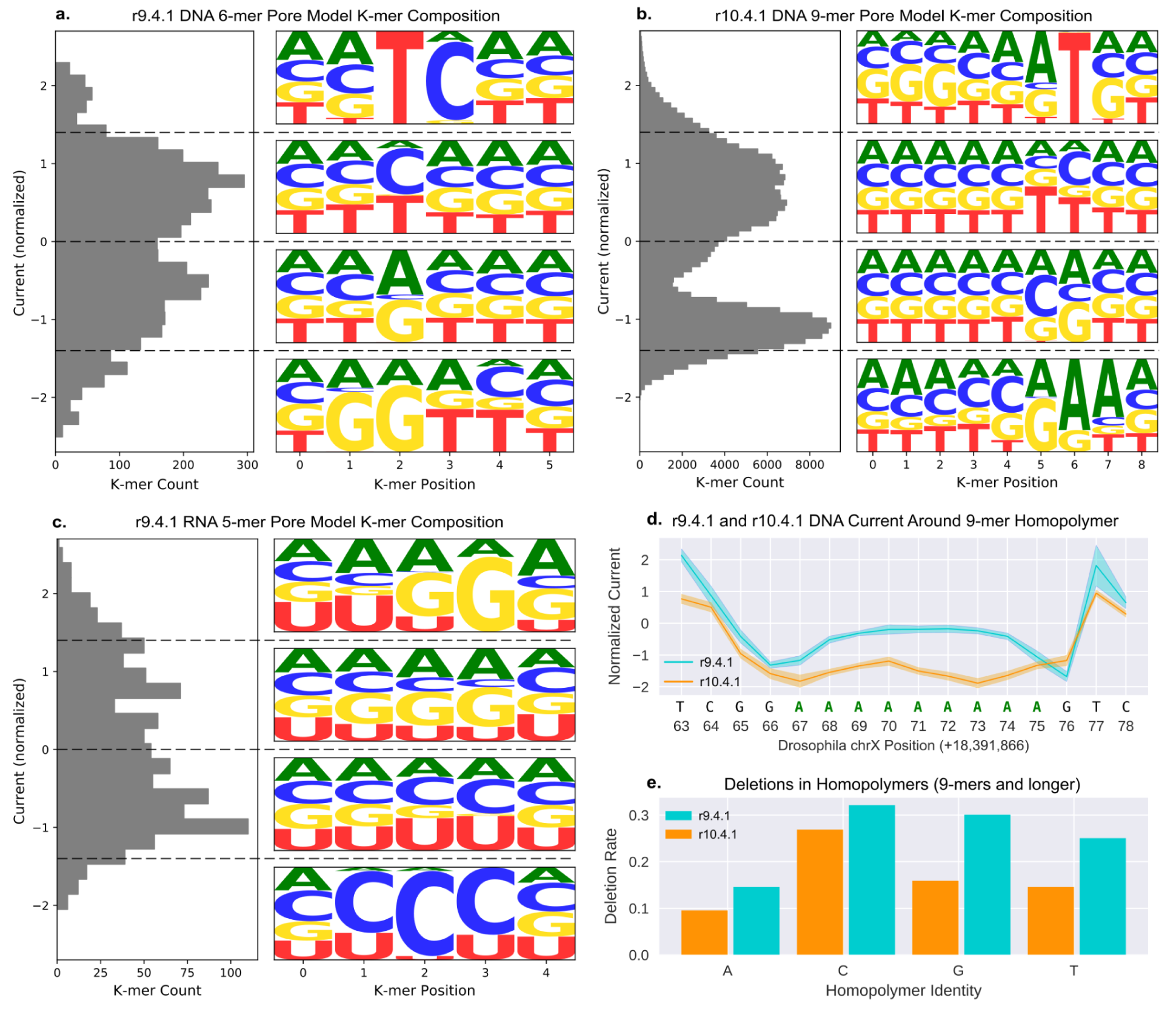
Current distribution and nucleotide composition of k-mers in Uncalled4 trained pore models. Plots represent **(a)** r9.4.1 DNA, **(b)** r10.4.1 DNA, and **(c)** r9.4.1 RNA (RNA002). ONT pore models are highly similar and produce nearly identical figures. **(d)** Mean and standard deviation of current surrounding a 9-mer adenine homopolymer in the *D. melanogaster* genome, based on Uncalled4 alignments of r9.4.1 and r10.4.1 DNA reads. **(e)** Fraction of basecalled reads containing a deletion within homopolymers of length nine or longer in the *D. melanogaster* X chromosome, computed using samtools mpileup.

### Pore model characteristics

To understand the nucleotide composition of pore models, we generated sequence logos of k-mers within different current ranges for each model (**Fig. 2a-c**). In both DNA models we find purines (A and G) at the central base enriched at low current levels and pyrimidines (C and T) enriched at high current levels. RNA has a weaker high-level relationship between nucleotide content and current, where we see a less consistently dominant effect at the central base. These high-level sequence logos are nearly identical for ONT’s pore models and the corresponding models trained by Uncalled4 (see below).

The reported intent of r10.4.1’s double reader head is better accuracy around homopolymers. Homopolymers longer than the span of the reader head register little-to-no change as the same bases repeatedly pass through the pore, generating a higher frequency of deletions (**Fig. 2d**). Supporting this intent, we find r10.4.1 has a lower frequency of deletions in homopolymers nine nucleotides or longer in the *D. melanogaster* genome compared to r9.4.1 (**Fig. 2e**). The deletion rate varies depending on which nucleotide the homopolymer is composed of, more so than the overall difference between r9.4.1 and r10.4.1, and is highest in cytosine with 26% of reads containing a deletion.

### Pore model training

Uncalled4 iteratively trains pore models by repeatedly aligning signal and averaging signal characteristics for each k-mer (see **Methods**). Beginning with DNA, we trained models to represent canonical nucleotides using PCR-amplified *Drosophila melanogaster* DNA, chosen as its genome contains every possible 11-mer, providing enough context to test different k-mer lengths without requiring more extensive amounts of sequencing. To train models without a prior pore model, we use the *ref-moves* used to guide our bcDTW algorithm. For r9.4.1 DNA we initialize the process with a single nucleotide (1-mer) model and expand the k-mer length every-other training iteration, alternating which side to append the new nucleotide, to generate a 6-mer model nearly identical to ONT’s proprietary model (Pearson’s r=0.9995) (**Supplemental Fig. 3a**). A similar process was used for r10.4.1, but we initialized with 4-mers and expanded to generate 9-mers, matching ONT’s model with slightly lower but strong Pearson’s correlation (r=0.9928) (**Supplemental Fig. 3b**).

We note a large group of outliers in comparing Uncalled4 and ONT’s r10.4.1 models with a consistent NNNNTVTTN motif (N = any base, V = not T). The current for these k-mers is inconsistent with all other k-mers with Ts in two central positions which are otherwise a strong predictor for current level (**Supplemental Fig. 3c**). The ONT pore model anomalies are further supported by comparing to ONT’s r10.4.1 260bps model, representing a deprecated option for sequencing at slower speed, which has no such outliers and higher similarity to Uncalled4’s 400bps model (r=0.9979) (**Supplemental Fig. 3d**). Finally, calling 5-methylcytosine CpG methylation (see below) in the CGNTVTT context yields a weaker secondary peak when using the ONT model, suggesting poorer alignments in this context (**Supplemental Fig 3e**). Taken together, this suggests that the ONT r10.4.1 400bps pore model is inaccurate in the TVTT context, highlighting the value of our fully reproducible pore model training method.

We also trained a r9.4.1 direct RNA (RNA002) model using *in vitro* transcribed (IVT) human HeLa cell line data, using a similar process as r9.4.1 DNA but with additional iterations-per-k-mer-length to adjust for additional noise. The Person’s correlation with the ONT model was lower than either DNA model but still strong (r=0.9804). Interestingly, we note a stronger similarity between our RNA002 model and the five central bases of ONT’s RNA004 model (r=0.9863), the latest RNA sequencing chemistry recently announced by ONT (**Supplemental Fig. 4**). These results indicate the current characteristics of RNA004 are highly similar to RNA002, so that Uncalled4 can be used with this model as these data become available.

### r10.4.1 5-methylcytosine DNA model training and detection

To explore the effect of DNA modifications on the r10.4.1 DNA sequencing, we sequenced *D. melanogaster* DNA treated with CpG methyltransferase M.SssI to broadly modify CpG sites with 5-methylcytosine (5mCpG), with an average per-site methylation rate of 88% estimated by the Guppy basecaller (v6.4.8, high-accuracy model). We trained a 9-mer model using the process described above and compared the current levels to the unmodified model, confirming that k-mers with CpG in the central position were the most divergent (**Fig. 3a**). In contrast, CpGs in the first five k-mer positions (secondary reader head) provide almost no information (**Supplemental Fig. 5a)**. K-mers with CpG adjacent to the central position exhibit some divergence depending on the base identity, consistent with the model substitution profiles (**Supplemental Fig. 5b-c**, Fig 1a).

Next, we tested Uncalled4’s ability to directly detect CpG methylation by comparing current levels between PCR and 5mCpG *D. melanogaster* r10.4 data using nonparametric Komologorov-Smirnov (KS) statistics. We included all CpG sites with a 100% modification rate estimated by Guppy and aggregated all KS statistics over a window surrounding each CpG site. We compared Uncalled4 with f5c, the only other aligner currently capable of r10.4.1 alignment. Both generated two peaks in KS statistics around CpG sites, with a strong peak upstream and a weaker peak downstream, consistent with the double reader head (**Fig. 3b**). Uncalled4 computed a higher primary peak than f5c, indicating a clearer signal of modification, and both aligners generated similar secondary peaks. Uncalled4 has generally higher KS statistics overall, consistent with less signal masking than f5c, but the relative difference around CpG sites remains higher for Uncalled4. The overall KS statistic trend is the same using ONT’s r10.4.1 400bps k-mer model or the Uncalled4 trained model, but as noted previously the Uncalled4 model performs better in CGNTVTT contexts with a stronger secondary peak (**Supplemental Fig. 3e**).

**Figure 3.**
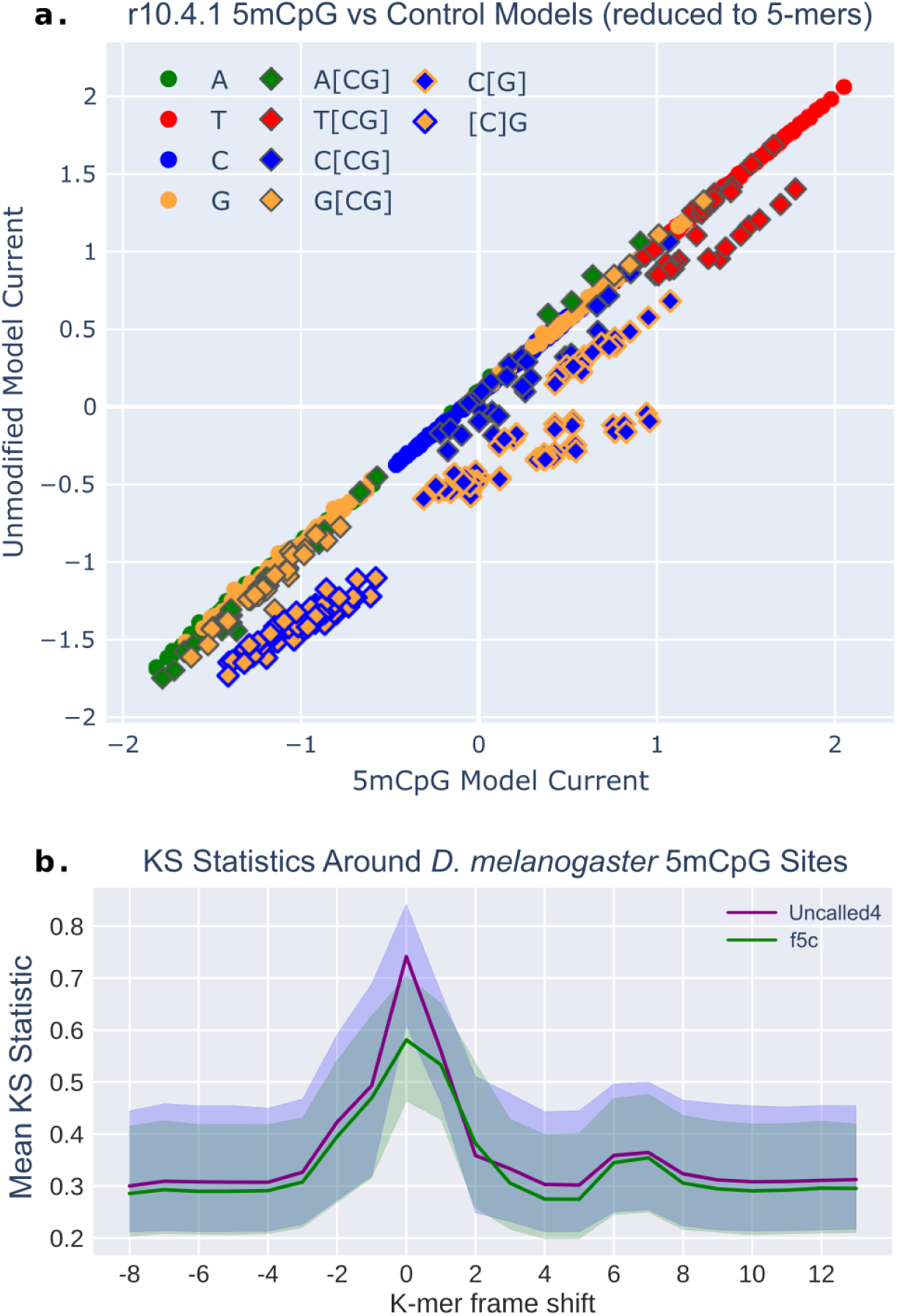
5mCpG signal characteristics. **(a)** Normalized current levels for Uncalled4 5-methylcytosine (x-axis) and unmodified control (y-axis) r10.4.1 pore models, reduced to 4-mers by averaging k-mers sharing their last four bases. Each point is colored by the identity of the central base, with diamonds representing CpG containing k-mers. Outlined diamonds indicate k-mers with the modified cytosine at central position (C[G]) or one base upstream ([C]G). **(b)** Current-level KS statistic mean and interquartile ranges surround 5mCpG sites in the *D. melanogaster* X chromosome, computed from Uncalled4 and f5c r10.4.1 signal alignments using the ONT r10.4.1 400bps model.

### Comparative RNA modification detection

To compare the effectiveness of Uncalled4, Nanopolish, and Tombo in RNA modification detection, we begin with a comparative approach using KS statistics between modified and unmodified RNA. Uncalled4 has consistently higher AUROC and AUPRC in detecting a diverse set of 36 modifications in *Escherichia coli* ribosomal RNA, based on KS statistics between native and *in vitro* transcribed (IVT) RNA at 100x coverage (**Supplemental Figure 6**). We apply similar methods to detect m6A in human RNA, the most abundant RNA modification in humans. For this, we use two HEK293T cell line samples: wildtype (WT) and METTL3 m6A methyltransferase knockout (KO), where the KO sample has lower m6A levels at METTL3-sensitive sites. Accuracy was estimated using m6ACE-seq labels filtered for METTL3 sensitive sites [27]. Using KS statistics on sites with at least 20x coverage from primary transcriptome alignments, Uncalled4 outperforms Nanopolish and Tombo in AUPRC and AUROC, both in all contexts and limited to DRACH motifs (**Fig. 4a**, **Supplemental Fig. 7**). We performed a similar analysis with xPore [27], a comparative RNA modification detection method that uses Gaussian mixture models (GMMs), using Nanopolish and Uncalled4 alignments of the same data, again yielding higher AUPRC and AUROC for Uncalled4 (**Fig. 4a**, **Supplemental Fig. 7-8**).

Moving to a supervised m6A detection method, we input Uncalled4 and Nanopolish alignments into m6Anet [28], a neural network method designed for Nanopolish alignments which calls m6A modifications at DRACH sites on individual reads before aggregating them to the transcript-level. m6Anet does not require a secondary sample to compare against, but the neural network did necessitate re-training for Uncalled4 alignments (see **Methods**). To compare m6Anet with comparative methods, we used the difference in modification rate at each site between the WT and METTL3 KO samples multiplied by the WT modification probability as the differential modification score, yielding higher AUPRC and AUROC with Uncalled4 than all other comparative methods (**Fig. 4a, Supplemental Figure 7-8**). Interestingly, Nanopolish+m6Anet has a lower AUROC than Uncalled4+KS statistics, and has nearly the same AUPRC (only 0.02 higher), demonstrating that the alignment method has a larger impact than the downstream detection method in this context.

**Figure 4.**
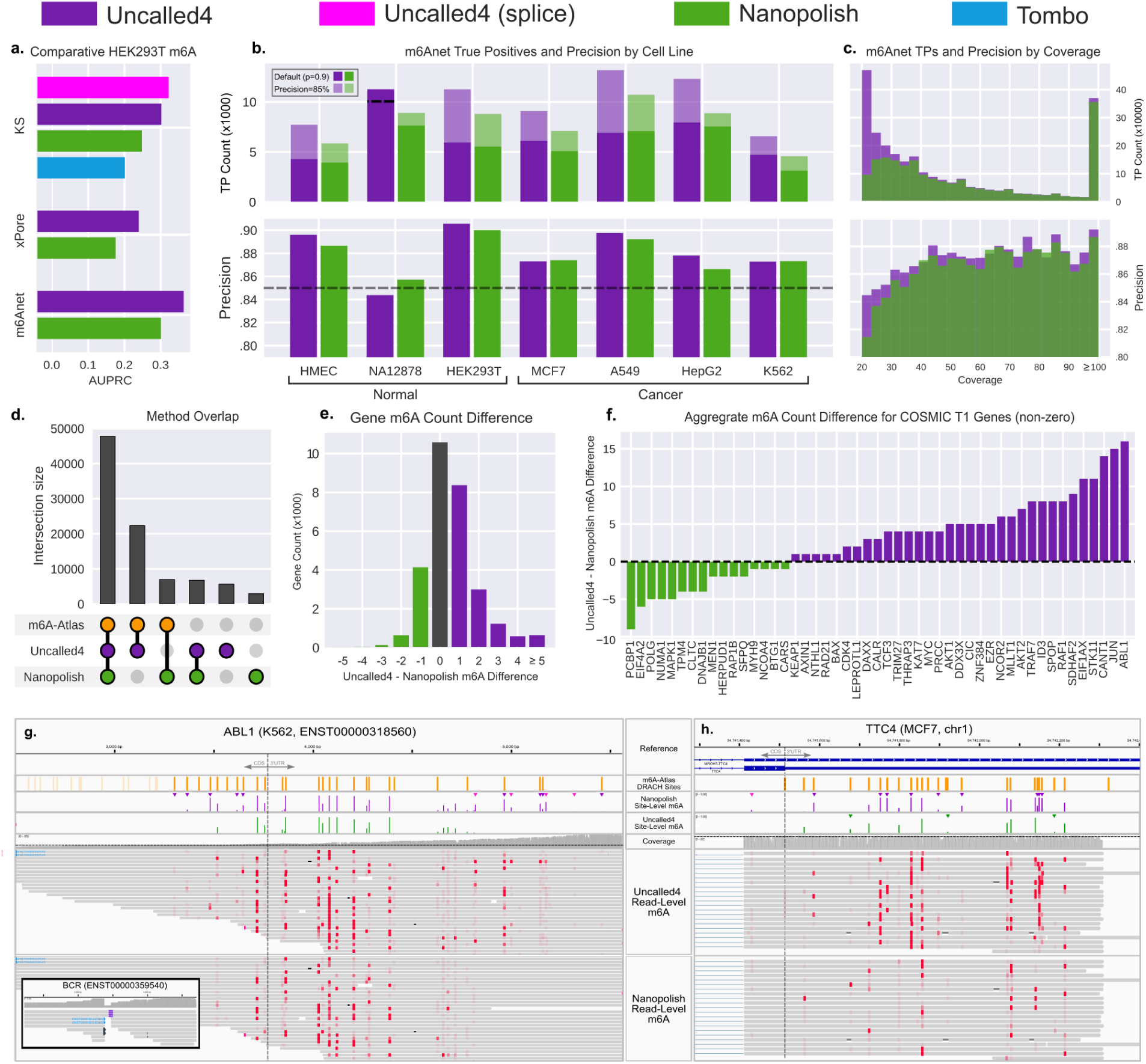
RNA modification detection. **(a)** Gene-level comparative m6A detection in DRACH contexts. “Uncalled4 (spliced)” (magenta) is based on spliced genome alignments, while all other use transcriptome alignments averaged to the gene-level. **(b)** Number of m6A sites found in each cell line which occur in the m6A-Atlas v2. Solid bars indicate the number of sites found with the default probability threshold 0.9, and shaded bars indicate the count at threshold where the precision is 85%. Uncalled4 with NA12878 has reduced recall at 85% precision, as indicated by dashed line. Precision with default probability threshold of 0.9. **(c)** Coverage distribution of true positive (TP) sites (top) and precision of sites within coverage bins. **(d)** Number of sites shared by Uncalled4, Nanopolish, and m6A-atlas v2 across all cell lines. **(e)** Difference in per-gene m6A count found by Uncalled4 and Nanopolish across all seven cell lines. **(f)** Difference in aggregated gene m6A count from Uncalled4 and Nanopolish alignments in COSMIC tier 1 genes with m6A modification found in every cell line (51 genes). **(g)** Transcript-level m6A calls in an ABL1 transcript alongside BCR fusion, and **(h)** gene-level m6A calls in the TTC4 gene.

While transcript-level modification calling is a unique strength of ONT direct RNA sequencing, gene-level modification calls may be more robust by combining information from multiple isoforms, especially when it is ambiguous from which transcript the read originated. Uncalled4 can perform genome alignment given spliced basecaller alignments, unlike Nanopolish and Tombo which only support transcriptome alignment for RNA. An alternative to direct genome alignment is mapping transcript-level calls to genome coordinates and averaging the probabilities (“t2g”, **Supplemental Fig. 9a**), a process that is built-in to xPore and has been applied to m6Anet. We include all multi-mapping reads for *t2g* mapping with Uncalled4 and Nanopolish to preserve ambiguously aligning reads from multi-isoform transcripts, finding that this increases AUPRC and AUROC compared to primary alignments (**Fig. 4b, Supplemental Fig. 9-10**). Note that this would not be appropriate for transcript-level analysis since the read aligner may clip alternatively spliced regions, and only primary alignments are included for Tombo as each read is fundamentally associated with a single alignment in its FAST5 format. Uncalled4 spliced genome alignment outperforms *t2g* mapping with KS statistics, and has higher AUPRC and AUROC than Nanopolish+m6Anet *t2g* mapping. Spliced genome alignments cannot be input to xPore or m6Anet as the tools cannot handle splicing or large chromosomes.

### m6A RNA across Cancerous and Healthy Human Cell Lines

We next compared the prevalence of m6A modifications using Uncalled4 and Nanopolish alignments with m6Anet in seven different human cell lines, consisting three normal and four cancer tissues (**Fig. 4b**). We used two replicates for most samples, except for HMEC where only one replicate was available, and HEK293T where three were used (**Supplemental Table 2**). We used m6A-Atlas v2 as our approximate set of m6A-positive sites, but note uneven representation across cell lines in m6A-Atlas, most notably HEK293T with over 100,000 gene-level sites, while others have no representation (NA12878, HMEC, K562) (**Supplemental Fig. 11a**). Nevertheless, the atlas provides a catalog of those bases that are commonly modified as a baseline for comparison. We first compared Uncalled4 and Nanopolish performance on a single wildtype HEK293T replicate using either HEK293T-specific m6A-Atlas labels or all m6A-Atlas labels regardless of cell type, in both cases yielding higher AUPRC and AUROC for Uncalled4 (**Supplemental Fig. 11b-c**). Limiting the m6A labels to those identified in HEK293T reduces the apparent performance for both aligners due to increased putative false positives, highlighting the limitations of m6A-Atlas as a ground truth as the epitranscriptome is highly dynamic and the m6A-Atlas is based on short-read assays with limited resolution. In either case, “recall” is likely underestimated since m6A-Atlas combines labels from many samples, and we do not expect consistent modification under all conditions. For all following analyses which include additional cell lines, we call each positive modification site a putative “true positive” (TP) if it occurs in any sample in the m6A-Atlas, acknowledging that this is an imperfect truth set and accuracy will vary between cell lines.

Beginning with transcript-level m6Anet calls on primary alignments, we find that Uncalled4 alignments yield consistently higher AUPRC and AUROC than Nanopolish on all samples (**Supplemental Table 3**). Uncalled4 also has generally higher recall and precision using the default probability cutoff of 0.9, with 18% more true positive m6A sites on average. The same patterns are observed when probabilities are averaged to the gene-level, again including all multi-mapping reads, with more sites found on the gene-level by both aligners in every sample and Uncalled4 again finding 18% more true positives on average (**Fig. 4b-c, Supplemental Table 4**). Precision is lowest for NA12878 using either aligner, even more so than the other samples absent from m6A-atlas (HMEC and K562), likely because NA12878 has the lowest data quality as measured by observed yield, basecaller pass/fail rate, and quality scores compared to other cell lines (**Supplemental Table 2**). To correct the variable precision estimates, we adjusted the probability thresholds for each sample such that each has equal precision of either 80%, 85%, or 90% (**Supplemental Table 3**). Uncalled4 finds 27.3% and 28.1% more TP sites on average at 80% and 85% precision respectively, but finds 0.1% fewer TP sites at 90% precision. The lesser performance at 90% precision is likely due to the unreliability of m6A-Atlas labels at this threshold, where notably Uncalled4 finds more TP sites in HEK293T, the most represented cell line in the m6A-Atlas. We therefore use the set of modifications found at 85% precision for further analysis, yielding higher recall than the default threshold in every cell line except NA12872 (**Fig. 4b**).

Uncalled4 finds disproportionately more m6A at sites at low coverage due to Nanopolish’s pervasive masking, which often reduces the apparent coverage to below 20x minimum set by m6Anet (**Fig. 4c**). Sites found by either aligner are generally enriched around stop codons and in the 3’UTR, consistent with previous findings (**Supplemental Fig 12a-b**). Most m6A sites that are only found by Uncalled4 are in the m6A-Atlas, and most sites not present in the m6A-Atlas are found by both Nanopolish and Uncalled4 (**Fig. 4d**). For both methods, approximately half of all sites were only identified in one sample, 30-32% of which are absent from the m6A-Atlas. The total number of m6A sites generally decreases with the number of supporting samples, as does the putative false positive rate, with the notable exception of sites shared by all seven samples, which is greater than those shared by only six samples (**Supplemental Fig 12c**). This set of m6A sites shared by all samples indicates transcripts that are broadly modified; for example the gene c-Myc has seven modifications found by Uncalled4 in all samples, where m6A has a well studied stabilizing effect on the c-Myc mRNA [19].

Aggregating the per-sample gene counts further, we compute the total number of modifications found across all samples for each gene by Uncalled4 and Nanopolish, revealing Uncalled4 broadly finds more modifications than Nanopolish (**Supplemental Table 4**). Specifically, we find more m6A in 66% of genes, fewer in 18% of genes, and an equal number in 16% of genes (**Fig. 4e**). To further explore differences in m6A count in genes across the healthy and cancerous cell lines, we focus our analysis on COSMIC Census tier 1 genes [29], which identifies genes with mutations implicated in cancer development, and specifically only those with m6A present in all samples (**Fig. 4f**). Among this subset, the gene with the largest increase in m6A count between Uncalled4 and Nanopolish is *ABL1*, an oncogene which fuses with *BCR* in chronic myeloid leukemia (CML). *ABL1* has been identified as a potential target of the ALKBH5 demethylase [30], and it has been observed that m6A contributes to aberrant translation in BCR-ABL1 positive CML cases [31]. Incidentally, we find multiple m6A-containing reads which support the BCR-ABL1 fusion in the CML K562 cell line, demonstrating the long-range information provided by Nanopore sequencing (**Fig 4g**). The gene with the next-highest increase in m6A count is the oncogene *JUN*, which is a known target of the METTL3 methyltransferase and its translation is promoted by m6A modification [21]. Several other genes in this subset are known to be transcriptionally destabilized by m6A: *STK11* [17], *ID3* [32], *AKT1*, *AKT2* [18], and *NCOR2* [30]. Others are known to be stabilized by m6A, such as *c-MYC* [19] and *THRAP3* [20].

Furthermore, among the top ten genes ranked by increase in m6A sites with Uncalled4 is *TTC4*, with a total 60 sites identified by Uncalled4 and 35 by Nanopolish across all cell lines, notably none of which are in the m6A atlas (**Supplemental Table 4**). However, closer inspection revealed m6A Atlas assigned all *TTC4* labels to the *MROH7*-*TTC4* readthrough transcript, which entirely contains *TTC4* (**Fig 4h**). Most reads which align to *MROH7*-*TTC4* also multimap to *TTC4* and *TTC4* m6A modification has been implicated in lung sepsis response in mice [33], suggesting that m6A Atlas has mislabeled which gene the genomic coordinates should correspond to. *TTC4* contains a 34 amino acid tetratricopeptide repeat, which makes short-read alignment less reliable and may contribute to inaccuracies in its m6A-Atlas labels.

## Discussion

Raw nanopore signal is information-rich, encoding much more than the four canonical bases obtained from standard basecalling. Signal-to-nucleotide alignment is a critical step in extracting this information, but the process is error-prone and few standards exist for comparing alignment methods. Uncalled4 features a rapid and highly accurate alignment algorithm guided by basecaller metadata, a compressed and indexed BAM-based signal alignment file format, and analyses to facilitate comparisons between signal alignment methods. Uncalled4’s pore model training method is fully reproducible, requires no prior k-mer based model, and reveals potential errors in ONT’s official r10.4.1 DNA model. Accurate signal alignment enables more sensitive DNA and RNA modification detection than comparable signal aligners, enabling it to find substantially more RNA m6A sites in several disease-relevant genes using m6Anet in healthy and cancer human cell lines compared to Nanopolish.

A major benefit of epigenetic profiling with long-reads is that the genetic identity is maintained, in contrast to short-read methods which involve base substitutions or read truncation, making it possible to comprehensively measure single-nucleotide, structural, and epigenetic variation in one assay. In principle, these methods could be applied to a wide variety of samples with publically available nanopore sequencing data, however the raw signal required to identify modifications is often not made available, mostly due to large file sizes and lack of database support. Uncalled4’s BAM format efficiently provides the statistics required by most signal-based detection methods in a widely supported format. A similar BAM tag was recently introduced by Squiguliser [34], a nanopore signal alignment visualizer in-part inspired by an early version of Uncalled4, however this only stores signal coordinates and not the current-level data required for modification detection. The use of efficient and indexable data representations will become even more critical as long-read sequencing becomes more widely adopted. In addition to widespread clinical sequencing with long reads, the Human Pangenome Reference Consortium is using both ONT and PacBio sequencing to assemble a haplotype-resolved human pangenome [35]. Long-read pangenomes present the opportunity and challenge of pan-epigenomic analysis, complicated by every cell having a potentially unique and dynamic epigenome, and multiple types of nucleotide modification present across species. Uncalled4 provides a step toward scaling such analyses as more data becomes available.

Even more daunting than pan-epigenomics, a pan-epitranscriptomic catalog would need to account for the underlying dynamic nature of the transcriptome and a much wider array of RNA modifications. The most well studied RNA modifications play an important role in RNA stability, mRNA splicing, mRNA export, translation efficiency, and several other important roles [36]. Lack of training data is now the major factor preventing identification of most of the over 150 known RNA modifications. Certain modifications may also generate minute changes in signal, meaning accurate signal alignment is necessary to reveal these subtle changes. For modifications which can be detected but not identified for lack of accurate labels, Uncalled4’s visualizations (**Fig. 1**) and analyses can serve as useful exploratory tools.

Our work showed that aggregating transcript-level m6A calls to the gene-level is a straightforward approach to improve accuracy, however this eliminates potentially interesting transcript-specific results. Detailed exploration of transcript-level modifications cannot rely solely on labels from short-read assays, which generally do not provide transcript-level specificity. Long-read methods must also be improved to accurately assign reads to transcripts in multi-isoform genes. Conventional transcriptome alignment often maps reads to incorrect transcripts by trimming alternatively spliced regions, or fails to include all potential mappings of fragmented reads, as shown here and in previous work [37]. Spliced genome alignment is an alternative approach which avoids isoform alignment ambiguity, but fully interpreting genome alignments would require mapping and disambiguating reads from the genome to transcriptome, similar to reference-guided transcriptome assembly [38]. If transcript-level modifications could be accurately identified, such methods could be applied to allele-specific modifications, similar to recent work in conventional transcriptomics [2]. Long-reads are also well-suited for characterizing unannotated transcripts, or non-canonical transcripts generated by structural variation or circular RNAs, the latter of which are associated with RNA modifications like m6A [39].

We have presented a toolkit for nanopore signal alignment and analysis, focusing on applications in nucleotide modification detection. Signal alignment is useful in other applications, for example in several recent rapid signal mappers designed for targeted sequencing [7,8,40,41]. Uncalled4 could be useful in optimization of such approaches, and the Python module is already used by Sigmoni for basic signal processing [40]. Uncalled4 will also be valuable in understanding future updates to nanopore sequencing chemistry, and will aid other signal-based methods in adapting to those changes.

## Acknowledgments

We would like to thank Jennifer A. Urban for the extraction of *D*. *melanogaste*r genomic DNA and helpful discussion. We would like to thank Ben Langmead, Omar Ahmed, Mohsen Zakeri, Mihaela Pertea, and Hasindu Gamaarachchi for their helpful discussions. This work was supported, in part, by NIH awards U01CA253481 and OT2OD002751(to M.C.S.), NSF award IOS-2216612 (to M.C.S.), NHGRI HG010538 and HG009190 (to W.T.), The Lustgarten Foundation award 90101412 (to M.C.S.), and the Commonwealth Foundation (to M.C.S.). Part of this research project was conducted using computational resources at the Maryland Advanced Research Computing Center (MARCC). WT has two patents (8,748,091 and 8,394,584) licensed to Oxford Nanopore. SK has received travel funding from Oxford Nanopore.

## Methods

### Uncalled4 Overview

Uncalled4 is a Python module and command line utility, with many computationally intensive subroutines implemented in C++ with Python bindings provided via PyBind11. The command line functionalities are split into several subcommands (**Fig. 1f**):

- *align* implements the basecaller-guided signal alignment algorithm, which outputs a BAM file by default.
- *convert* converts between signal alignment formats, where BAM and eventalign formats support input and output, Tombo FAST5s only support input, and m6Anet and TSV files only support output.
- *train* iteratively applies the alignment algorithm to train pore models and outputs a directory with k-mer models produced in each iteration.

The remaining commands are divided into analysis and visualization. The analysis commands are *refstats*, which outputs reference-level statistics (e.g. KS statistics), *readstats,* which outputs read-level statistics (e.g. mean normalized model difference), and *compare*, which compares two BAM files containing the same set of reads aligned using difference methods (e.g. Uncalled4 and Nanopolish). Visualization commands display interactive Plotly visualization, either as HTML files exportable to SVG or PNG, or as web browser sessions. *dotplot* displays one or more alignments of a signal read (**Fig. 1b-c**), *trackplot* displays one or more alignment tracks of many reads aligned to a reference region (**Fig. 1d**), and the *browser* command runs a local Dash web server which displays an interactive alignment track which can be clicked to display summary statistics, a dotplot, and per-reference statistics distributions (**Fig. 1e**).

The *align* command is described in the following “Signal Preprocessing” and “Basecaller-Guided Dynamic Time Warping” sections, *convert* in the “Alignment Encoding and Formats” section, *train* in “Pore Model Training”, analysis commands in “Analysis of Signal Alignments”, and visualization commands in “Visualizations”.

### Signal Preprocessing

Prior to alignment, the raw nanopore electrical signal must be preprocessed to reduce noise and correct for systematic bias in the current levels. First, the individual sensor readings (raw samples) are segmented into “events” using the same algorithm as UNCALLED [8], which uses rolling t-tests to group samples with similar current levels. This groups signal representing the same nucleotides, although variable sequencing speeds result in frequent “stays” (consecutive events representing the same k-mer, ∼50% of events) and less frequent “skips” (an event representing multiple k-mers, ∼1-5% of events). These events are stored with their sample start, length (proportional to dwell time), current mean, and current standard deviation. Event detection parameters are chosen depending on the sequencing chemistry, where RNA uses longer t-test window lengths than DNA to adjust for the slower sequencing speed.

After event detection, the event current means are iteratively normalized to correct for systematic deviation from the pore model. The most basic form of normalization is the method-of-moments, where the current levels are linearly scaled such that their overall mean and standard deviation match that of the pore model. Uncalled4 initially uses reference-guided method-of-moments, which uses the expected mean and standard deviation of current levels for the reference k-mers at the coordinates determined from the basecaller alignments. This method works better for low-complexity sequences compared to standard method-of-moments, which assumes random sequence content. Finally, by default Uncalled4 performs two iterations of alignment, and in the second iteration it uses linear regression between the aligned events and the expected current of reference k-mers from the first iteration. This second iteration is not performed in the *train* subcommand by default to avoid over-fitting to an error-prone model. The Theil-Sen estimator, a nonparametric regression algorithm used by Tombo, was also tested, but this was found to be less accurate and slower than simple linear regression.

### Basecaller-Guided Dynamic Time Warping

Uncalled4 uses Dynamic Time Warping (DTW) to align preprocessed signal to a reference sequence guided by basecalled read alignments. DTW is a widely used dynamic programming (DP) algorithm which has previously been applied to nanopore sequencing by Tombo and others [24,42]. Nanopolish uses a Hidden Markov Model (HMM) for alignment, which uses more complex transition probabilities which are trained on real data, but is otherwise similar to DTW in time and space complexity. The most basic form of DTW has O(NxM) complexity, where N is the number of read events and M is the number of reference k-mers. This can be improved using banded alignment, where the DP matrix is only filled in along the diagonal where the optimal alignment path is usually found. Tombo, Nanopolish, and f5c both use adaptive banded alignment, where the band position is adjusted as alignment progresses to always be centered on the currently most probable path.

Uncalled4 uses a dynamic banding algorithm similar to that described by f5c [23], but the band placement is chosen before alignment begins using the basecaller “moves” metadata. Basecallers such as Guppy and Dorado can optionally output “moves” which represent approximate alignments between the signal and the basecalled read. These moves have low-resolution (five samples for DNA, ten for RNA), and often deviate from the true alignment by one or more reference positions. Uncalled4 projects these basecaller moves into reference coordinates based on the basecalled alignment cigar string, then centers the DTW bands on these “reference moves” (**Supplemental Figure 1**). This allows Uncalled4 to use a much narrower bandwidth (25 by default) than Nanopolish or Tombo, making alignment faster and preventing alignments from straying too far from the truth. Insertions and deletions (indels) that are larger than the bandwidth would cause a discontinuity in the band placement, so these are “spliced” out of the read or reference respectively if they are above a threshold (10 by default) based on the *ref-moves* coordinates. Note Uncalled4 does not disrupt alignments over small indels, as these are a frequent basecaller error which can often be accurately aligned over. For deletions, the “splicing” generates k-mers that are not present in the reference, which many downstream tools cannot handle, so these are masked and not included in the output by default. However, this can be disabled with the “--unmask-splice” option.

Uncalled4 encodes alignments as per-reference-coordinate statistics (“layers”, see below), at a minimum consisting of raw sample coordinates assigned to each site. This is unlike Nanopolish eventalign, which outputs multiple consecutive events aligning to the same reference position. The first step of most modification detection algorithms is to average these statistics on the nucleotide-level, which is straightforward for the average current, but notably the current standard deviation cannot be accurately computed without re-analyzing the original raw signal. Uncalled4 outputs accurate per-nucleotide current standard deviations, which Tombo can also do via an optional flag. A consequence of this encoding is that “stays” and “skips” are not explicitly encoded by default. Skips can be identified by multiple consecutive reference positions having the same signal coordinates. Nanopolish masks skips and represents them as missing data. Uncalled4 penalizes skips in the DTW cost function (2x for standard alignment, 4x for pore model training), and they can be masked via the “--mask-skips [all|keep_best]” option, which removes either all grouped positions or all but the closest to the reference current. Skips are masked during pore model training to reduce alignment errors, however for modification detection we found that simply assigning the same values for each skipped position resulted in much higher recall with little change in precision. This reflects that modifications inherently disrupt signal alignment by deviating from the expected current, sometimes resulting in skips, and so masking skips removes useful information. This effect was most significant in RNA, where the motor speed is less consistent and skips may be caused by motor “slippage”. Stay and skip rate can be computed by including the command “uncalled4 align --count-events –tsv-out …”, which includes the number of events aligned to each reference position. Counts greater than one indicate stays, while fractional counts indicate the inverse of the number of skipped positions (e.g. 0.5 indicates two positions, 0.25 means two positions).

### Alignment Encoding and Formats

Uncalled4 represents each signal alignment as a set of per-reference-position statistics called alignment “layers”. All alignments must include the “length” layer, which indicates how many raw samples were aligned to each reference position, and usually include the current mean and standard deviation (“current” and “current_sd”). The current statistics are omitted for *ref-moves* due to their inaccuracy, and Tombo does not compute “current_sd” by default. Additional layers can be derived from the base layers and/or the reference sequence, such as “seq.kmer” (reference k-mer), “dtw.model_diff” (absolute difference between the read current and model current), or “mvcmp.dist” (distance from ref-moves) (**Supplemental Table 5**). One minor limitation of this reference-oriented encoding is that Uncalled4 cannot output event-level statistics, but rather averages over multiple events aligned to the same position. Tombo stores alignments in a similar manner, while Nanopolish outputs per-event statistics. Most modification detection tools simply average these statistics over reference coordinates, and in doing so cannot accurately compute the true per-base current standard deviation without re-querying the raw signal file (i.e. FAST5, SLOW5, or POD5). Uncalled4 computes the true current standard deviation at each position, and can optionally output the number of events aligned to each position.

Uncalled4 primarily stores signal alignments in BAM tags, alongside the conventional basecalled alignments that were used to guide bcDTW. This format differs from Nanopolish basecalled read and alignment paths are fully preserved, and unlike f5c’s similar format, Uncalled4 includes current means and standard deviations required for modification detection. This is accomplished in a space-efficient manner by storing the alignment layers in 16-bit integers. Raw signal coordinates (“us:” tag) are encoded with positive values indicating the number of aligned samples at each consecutive reference position, negative values indicating masked signal, and zeros indicating “skip” events (grouped with previous position). Most positions fit within 16-bit integers, and for the few outliers that cannot, we reserve the maximum value of 2∧16-1 to be grouped with the subsequent length entry. Reference coordinates (“ur”: tag) are encoded as a series of “start” and “stop” values indicating stretches of continuous alignment, with breaks caused by introns or deletions greater than “--del-max” (10nt by default). The total span of the reference coordinate blocks should be equal to the number of non-zero elements in the “us:” tag, and the sum of the absolute values of “us:” should equal the length of the raw signal. Current means (“uc:” tag) and standard deviations (“ud:” tag) are represented as 16-bit fixed-precision floating point values corresponding to normalized current levels ranging from -5.0 to 5.0 by default, representing a range of five standard deviations from the mean. Masked positions are assigned a “null” value equal to -2^16^-1. Normalized units can be converted to picoamps, or whichever units are defined by the pore model, using parameters stored in JSON format in the BAM header. This JSON header stores additional information on the tag labels, fixed-point scaling factors, reference and raw signal paths, and other pore model metadata. The addition of the signal alignment tags increases the BAM file size by 2-4 fold, but is still several times smaller than the Nanopolish and Tombo formats (**Table 1**).

In addition to the BAM format, Uncalled4 supports two text-based output formats: “eventalign” and “TSV”. Eventalign is based on Nanopolish’s default tab-delimited output format, and is mainly included for compatibility with modification detection tools like xPore and m6Anet. “TSV” is a customizable tab-delimited format, which can include any of the alignment layers or comparison statistics (see below). These formats can be written directly by the “uncalled4 align” command, or can be derived from an Uncalled4 BAM file via the “uncalled4 convert” command with the “--bam-in” option. “uncalled4 convert” can also convert Nanopolish eventalign files or Tombo FAST5 files into the BAM format, which is necessary for analysis and visualization of these alignments by Uncalled4.

Finally, to expedite m6Anet analysis and demonstrate the utility of the BAM alignment format, Uncalled4 includes a conversion function from a sorted BAM file to the m6Anet “dataprep” format that collects signal features in a per-reference-coordinate JSON format. This can also be accomplished by first converting the BAM file to “eventalign” format and using “m6anet dataprep”, however we found conversion from eventalign format was by far the largest bottleneck in m6Anet analysis. The sorted BAM format enables conversion in a single linear read of the file, making conversion many times faster than the random parallel file access required to convert from eventalign, especially on a shared compute cluster where parallel disk access is slow.

### Analysis of Signal Alignments

Uncalled4 can perform analysis on any signal alignments in the BAM format, which can be divided into reference-level (*refstats* command), read-level (*readstats* command), and read-base-level (*convert* and *compare*). Reference-level analysis includes simple summary statistics like mean and standard deviation of current levels and dwell times, or comparative statistics like KS statistics between two samples. Similarly, read-level analysis outputs summary statistics of layers over entire reads, or segments of reads defined by reference coordinates. Basic and derived layers of individual reads at each reference coordinate can be output in TSV format via the *convert* command. If basecaller *moves* are included in the BAM file, this can include *ref-moves* distance metrics (described below).

Uncalled4 can compare two different alignment methods applied to the same set of reads by inputting two sorted signal alignment BAM files to the *compare* command, producing a table of per-reference-coordinate *Signal Jaccard Distances* and *Signal-to-Reference Distances.* These can also be visualized via the *dotplot* command (**Fig. 1c**). *Signal Jaccard Distance*, the inverse of the Jaccard similarity, measures the degree of overlap between raw samples aligned to each reference coordinate: 1 - (A U B) / (A ∩ B), where A and B are the sets of raw samples aligned to each reference coordinate. This varies between 0 and 1, where 1 indicates no overlap and 0 indicates perfect overlap. *Signal-to-Reference Distance* measures the average number of reference bases between the raw samples aligned to each coordinate. This is computed reciprocally for each method, computing the nucleotide distance for each raw sample aligned to each reference coordinate, and then averaged between the two methods. These metrics can also be computed between a signal alignment method and the *ref-moves* used to guide Uncalled4 alignment, via the *convert* command or in any of the visualizations. In this case, *signal-to-reference distance* is *not* computed reciprocally, instead only averaging over the alignment method raw samples and not the low-resolution *ref-moves*.

### Visualizations

All visualizations produced directly by Uncalled4 are implemented in Python using Plotly, which produces interactive web-browser-based plots that were exported to SVG format. The three main alignment visualizations (trackplot, dotplot, and refplot) and also integrated into an interactive signal genome browser using Dash, a local web server designed for Plotly. Pore model profiles (**Fig 1a**) are generated using pore models only by computing the absolute change in current generated substituting each base for each other base at every possible k-mer. This is efficiently implemented in Python and C++ using the “buffer protocol”, which allows for vectorization of k-mer operations. Some figures were also generated by the Python matplotlib library, in cases where reproducible interactivity is not necessary.

IGV visualizations were generated by encoding the per-read modifications defined m6Anet’s “data.indiv_proba.csv” file into BAM modification tags. The reference coordinates were translated to read coordinates using the cigar string, and positions where an “A” was not present in the read sequence were excluded. Site-level probabilities were multiplied by a constant factor (4) for visualization purposes. The IGV screenshots were exported to the SVG format and edited in Inkscape for clarity.

### Pore Model Training

Pore model training is an iterative process, where in each iteration reads are aligned until every k-mer is represented a minimum number of times (500 by default), after which summary statistics (median and standard deviation) of signal characteristics (current mean, current standard deviation, and dwell time) are recorded and used as the pore model for the next iteration. In each training iteration, only positions with low *signal-to-reference distance* to *ref-moves* are included (*mvcmp.dist* <= 1, by default), which eliminates many alignment errors. Only one normalization iteration is applied during training, since linear regression is sensitive to outliers that are frequent in early training iterations. A higher skip penalty is also used for pore model training, and skipped positions are masked to further reduce alignment errors (equivalent to “uncalled4 align --skip-cost 4 –mask-skips keep_best”). This process requires an initial “draft” model to use in the first iteration. This draft model could be a canonical nucleotide model, for example with the goal of re-training it for modified nucleotides. We also developed a *de novo* initialization method, not requiring a prior k-mer pore model beforehand, using the *ref-moves* used in Uncalled4’s bcDTW algorithm. The *ref-moves* can be treated as standard signal alignments, although they are frequently one or two bases from their true position. To mitigate these inaccuracies, we began training using a short k-mer length to average-out the initial errors (1-mers for r9.4, 4-mers for r10.4) and increased the k-mer length every N training iterations (two iterations for DNA, three for RNA) until the desired k-mer length was reached.

The Uncalled4 *train* subcommand runs the training procedure for a specified k-mer length and number of iterations. It outputs a directory with the pore model for each iteration in a binary NumPy format, along with indexed alignment statistics used to generate each model. To progressively increase the k-mer length, the command was run once per-k-mer-length, each time using the last pore model output in the previous iteration as the new initialization model. This training procedure is flexible, allowing for alternate initialization methods or k-mer expanding methods to be tested. We evaluated the effectiveness of each training procedure based on the Pearson’s correlation coefficient between the trained model and ONT’s pore models. Intermediate models with shorter k-mer lengths can also be evaluated by “reducing” ONT’s models by averaging the values of k-mers which share central bases, implemented in Uncalled4’s “PoreModel.reduce” method.

The *align* subcommand will attempt to automatically detect the appropriate pore model to use based on metadata in the raw signal FAST5/POD5/SLOW5 files. If this can not be detected, the user can specify a preset pore model (“dna_r10.4.1_400bps_9mer”, “dna_r9.4.1_400bps_6mer”, or “rna_r9.4.1_70bps_5mer”) using “--pore-model” flag or by defining the “--flowcell” and “--kit” used for basecalling, or a custom pore model can be provided. The default r9.4.1 DNA and RNA pore models are provided by ONT (https://github.com/nanoporetech/kmer_models), while for r10.4.1 DNA we use the Uncalled4 model trained on unmodified *D. melanogaster* data, as described above.

### Modification Detection, Training, and Assessment

Comparative KS statistics were computed by the Uncalled4 *refstats* command: “uncalled4 refstats current.mean ks …”. For Tombo alignments, we compared the output to the Tombo KS statistic output, with highly similar results. xPore was run on Nanopolish using recommended parameters, and on Uncalled4 via conversion to the eventalign format.

m6Anet was trained for Uncalled4 alignments on the HCT116 cell line from the Singapore Nanopore Expression Project (replicate 3) [43], which was originally used for the default m6Anet model [28]. This data was re-basecalled using Guppy v6.4.8, and we also re-trained Nanopolish to assess the effect of re-basecalling. m6A labels obtained from [28] were provided in transcript coordinates, with sites divided by-gene into “train”, “test”, and “validation” sites. The recommended training procedure only included primary transcriptome alignments, and we noted that many reads aligned to a different transcript from the same gene than was listed in the training data. We therefore mapped the training data to gene-level coordinates, then back to transcript-level using the transcripts present in the re-basecalled data, maintaining the same “train”, “test” and “validation” gene assignments.

Precision-recall and receiver operator characteristic (ROC) curves were visualized and the corresponding area under the curve (AUC) were computed using Scipy. Both these metrics measure recall, also known as true positive rate (TPR), defined as TP/(TP+FN) (TP is true positive count, FN is false negative count). Accurate estimation of FN is complicated by pre-filtering performed by tools like xPore and m6Anet, where many sites are not assigned a probability and excluded from the output. We found many of these sites are actually modified, and so not counting these decreases the FN count and thus falsely increases the recall (**Supplemental Figure 8**). We used two strategies to compensate for this. For the transcript-level HEK293T results, we computed coverage from minimap2 alignments of basecalled reads, only considering read endpoints and not internal deletions, and included all sufficiently covered (20x) sites by filling in a probability of zero. For the gene-level results, different alignment strategies cover different sites, and so we simply took the union of all sites covered by each tool and filled missing values with probability zero. Many sites with probability zero can generate a large “jump” at the end of precision-recall and ROC curves, making the AUC estimate less informative. For precision-recall, we use the “average precision” definition of AUPRC, which essentially treats the curve as a stepwise function rather than using the trapezoidal rule for area calculation, which is more robust to skewed datasets. There is no comparable alternative for AUROC, but visual inspection suggests the overall trends would be the same regardless.

### Data Processing

All nanopore read data was basecalled with Guppy v6.4.8 using high-accuracy models wit the “--moves_out” option, with the exception of Tombo alignments where an earlier version of Guppy (v6.0.1) which supported output of basecalled FAST5 files required for Tombo. 5mCpG calling was also performed using the Guppy 5mC high-accuracy model. Reads were aligned using Guppy’s builtin minimap2 alignment option, which encodes the “moves” basecaller metadata in primary alignment tags. We also wrote a Python script which copies these tags into supplemental and secondary alignments for efficient alignment with Uncalled4. To re-align reads while preserving basecaller metadata, for example to compare spliced genome alignments with transcriptome alignment, we converted the primary BAM alignments to FASTQ with the relevant tags in the header using the command “samtools fastq -T mv,ts”, then aligned using minimap2 (v2.16) with the “-y” option to copy tags from the headers.

The *D. melanogaster* data was aligned to the *D. melanogaster* ISO1 release 6 reference genome (RefSeq GCF_000001215.4). The *E. coli* rRNA data was aligned to the 16S and 23S transcripts from the *E. coli* transcriptome (GenBank NC_000913.3), with modification labels obtained from [44]. All human datasets were aligned to a transcriptome derived from GRCh38 Ensembl annotations version 91 obtained from [27], or directly aligned to GRCh38 for spliced genome alignments (GCF_000001405.26). HEK293T METTL3-sensitive m6A labels were also obtained from [27]. HCT116 m6A training labels were obtained from [45]. All other m6A labels are from the m6A-Atlas v2 [16] (accessed May 12th 2023).

Uncalled4 signal alignments were compared with Nanopolish (v0.13.3), Tombo (v1.5.1), and f5c (v1.3). Nanopolish and f5c were run using the “--scale-events --signal-index” options, which are required for Uncalled4, xPore, and m6Anet. Timing was measured using a single CPU thread, and f5c additionally utilized a Nvidia Quadro P5000 GPU with default parameters. Komolgorov-Smirnov (KS) statistics were primarily computed with Uncalled4, and produced similar results to Tombo’s builtin KS statistic output, with minor differences attributed to read filtering and rounding error. We also compared Uncalled4 and Nanopolish RNA modification detection performance using xPore (v2.1) and m6Anet (v2.0.2). m6Anet was re-trained using Uncalled4 and Nanopolish alignments on re-basecalled HCT116 cell line data and labels originally used for m6Anet. GNU parallel was also used to efficiently run tasks in parallel.

### DNA extraction

Genomic DNA was extracted from 15 newly eclosed *D*. *melanogaster* males of the Oregon R strain (Bloomington stock center #5, RRID: BDSC_5). After selection, males were immobilized by freezing at -80°C for 5 minutes. Next, the flies were crushed with a pipette tip in 200 uL of Buffer A (100 mM Tris-HCl, pH 7.5, 100 mM EDTA, 100 mM NaCl, 0.5% SDS). This was followed by a 30-minute incubation at 65°C. After incubation, 400 uL of KOAc:LiCl (prepared by combining 1 part 5 M KOAc with 2 parts 6 M LiCl) was added and the mixture was allowed to precipitate on ice for 10 minutes and then the precipitate was pelleted at room temperature at 14,000 RPM for 15 minutes. The supernatant containing nucleic acids was transferred to a clean microcentrifuge tube and isopropanol was added at a ratio of 600 uL per 1 mL supernatant. The DNA was then precipitated by centrifuging at 14,000 RPM for 15 minutes at room temperature. Afterward, the supernatant was removed, and the pellet was washed with 1 mL of cold ethanol (70-75%). The pellet was then centrifuged again for 5 minutes before removing the ethanol wash. After air drying, the pellet was resuspended in ultra-pure water.

### Genomic DNA shearing and amplification

*D. melanogaster* genomic DNA (∼500 ng) was diluted into a total volume of 49 uL ultra-pure water. In order to shear the genomic DNA to 8 Kb fragments, DNA was transferred to a g-Tube (Covaris, 520079) and centrifuged at room temperature for 1 minute at 6000 RPM. The g-Tube was then inverted and centrifuged again at room temperature for 1 minute at 6000 RPM. Centrifugation was carried out on an Eppendorf Centrifuge 5425 (Eppendorf, 5405000646).

Sheared DNA was then amplified using the Oxford Nanopore Technologies (ONT) protocol for low input PCR (low-input-genomic-dna-with-pcr-sqk-lsk110-LWP_9117_v110_revJ_10Nov2020-minion) . First, the sheared DNA was mixed with NEBNext Ultra II End Prep Enzyme Mix and Reaction Buffer (NEB, E7180S) and incubated at 20°C for 5 minutes followed by 65°C for 5 minutes to repair fragment ends. DNA was then purified using 1x AMPure XP beads (Beckman Coulter, A63881) along with 70% ethanol. DNA was eluted in 31 uL nuclease-free water and quantified with the Qubit broad range dsDNA assay (ThermoFisher Scientific, Q32850). Next, end-prepped fragments were mixed with PCR adapters (ONT, EXP-PCA001) and Blunt/TA Ligase Master Mix (NEB, M0367S) and incubated for 15 minutes at room temperature. DNA was then purified using 0.4x AMPure XP beads and 70% ethanol. DNA was eluted in 26 uL nuclease-free water at room temperature and quantified with the Qubit broad range dsDNA assay. DNA was then diluted to 10 ng/uL in water.

Twelve PCRs were then performed by combining 20 ng diluted DNA, 46 uL water, 2 uL Primer Mix (ONT, EXP-PCA001), and 50 uL LongAmp Taq 2x master mix (NEB, M0287S). PCR cycling conditions were as follows:

| Temperature | Time | Number of cycles |
| --- | --- | --- |
| 95°C | 3 minutes | 1 |
| 95°C | 15 seconds | 18 |
| 62°C | 15 seconds | 18 |
| 65°C | 8 minutes | 18 |
| 65°C | 10 minutes | 1 |
| 4°C | hold | . |

Pairs of PCR reactions were then combined and DNA was purified using 0.4x AMPure XP beads and 70% ethanol. DNA was eluted at room temperature in 30 uL nuclease-free water and quantified with the Qubit broad range dsDNA assay. Fragment size was quantified using a Genomic DNA ScreenTape (Agilent, 5067-5365) on a TapeStation 4200 (Agilent, G2991BA). All DNA was then pooled and stored at -20°C until use.

### CpG 5-methylcytosine (5mC) labeling

Labeling of CpGs with 5mC to create a training dataset was performed similarly to previous work [22,46]. Two labeling reactions were set up as follows. Amplified DNA (4 ug) was combined with 40 uL water, 8 uL 10x NEB Buffer 2 (NEB, B7002S), 8 uL 1.6 mM S-adenosylmethionine (SAM)(NEB, B9003S) and 16 units of M.SssI (NEB, M0226S). Reactions were incubated for 4 hours at 37°C. After two hours of incubation, both 1.6 mM SAM (8 uL) and M.SssI (16 units) were added to the reactions to replenish enzyme activity. DNA was then purified using 0.8x AMPure XP beads along with 70% ethanol and eluted in 22 uL nuclease-free water. DNA was then quantified with the Qubit broad range dsDNA assay and fragment size was quantified using a Genomic DNA ScreenTape on a TapeStation 4200. DNA from the two reactions were then pooled and stored at -20°C until further use. A second round of M.SssI labeling on the pooled DNA was performed identically to the labeling reaction above. After final DNA purification and quantification, the DNA was stored at -20°C until sequencing.

### Nanopore library preparation

Four ONT sequencing runs were performed. Both unlabeled and labeled DNA were sequencing on r9.4.1 pores as well as r10.4.1 pores at 400 bases per second. Libraries for r9.4.1 pores were constructed using the LSK110 ligation sequencing kit (ONT, SQK-LSK110) and r10.4.1 pore libraries were constructed using the LSK114 ligation sequencing kit (ONT, SQK-LSK114). Both LSK110 (genomic-dna-by-ligation-sqk-lsk110-GDE_9108_v110_revV_10Nov2020-minion) and LSK114 (genomic-dna-by-ligation-sqk-lsk114-GDE_9161_v114_revG_29Jun2022-minion) have similar protocols so only one set of steps will be described below with notes on kit specific changes.

First, DNA fragments (1.25 ug) were mixed with NEBNext Ultra II End Prep Enzyme Mix and Reaction Buffer and incubated at 20°C for 5 minutes followed by 65°C for 5 minutes to repair fragment ends. DNA was then purified using 1x AMPure XP beads along with 70% ethanol, eluted in 61 uL nuclease-free water at room temperature, and quantified with the Qubit broad range dsDNA assay. End-prepped DNA was then mixed with Ligation Buffer (ONT, SQK-LSK110 and SQK-LSK114), NEBNext Quick T4 DNA ligase (NEB, E7180S), and either Adapter Mix F for r9.4.1 pores (ONT, SQK-LSK110) or Ligation Adapter for r10.4.1 pores (ONT, SQK-LSK114). Reactions were incubated for 15 minutes at room temperature. DNA was purified using 0.4x AMPure XP beads and Long Fragment Buffer (ONT, SQK-LSK110 and SQK-LSK114). DNA was eluted in 15 uL Elution Buffer (ONT, SQK-LSK110 and SQK-LSK114) at 37°C for 10 minutes. DNA was then quantified with the Qubit broad range dsDNA assay.

### Nanopore sequencing

R9.4.1 pore sequencing was performed using ∼40-50 fmol of library on r9.4.1 MinION flow cells (ONT, FLO-MIN106D). R10.4.1 pore sequencing was performed using ∼20 fmol of library on r10.4.1 MinION flow cells (ONT, FLO-MIN114) using either 260 or 400 bases per second mode. According to manufacturer recommendations, bovine serum albumin (Invitrogen, AM2616) was added to the sequencing flush buffer at a concentration of 0.2 mg/mL for all r10.4.1 flow cell sequencing runs. All sequencing runs were performed on a GridION Mk1 sequencing device (ONT, GRD-MK1) and run for 72 hours.

## Code and Data Availability

Uncalled4 is available open-source at github.com/skovaka/uncalled4. The *D. melanogaster* ONT sequencing data described above is deposited on the sequence read archive (SRA) bioproject PRJNA1082764. Trained pore models and the Uncalled4 m6Anet model are available on Figshare (https://doi.org/10.6084/m9.figshare.25336195.v1).

All other datasets were obtained from publicly available sources. The *E. coli* rRNA data was obtained from [44] (SRA bioproject PRJNA634693). The IVT HeLa cell direct RNA sequencing data used to train the RNA002 model was obtained from [47] (SRR23950400). HEK293T wildtype and METTL3 knockouts was obtained from [27] (PRJEB40872). NA12878 data was obtained from [48] (https://github.com/nanopore-wgs-consortium/NA12878). HMEC data was obtained from [49] (GEO accession GSE132971). All other human cell line data is from the Singapore Nanopore Expression Project (PRJEB40872, **Supplemental Table 2**).

