## Supplemental Figures for "Uncalled4 improves nanopore DNA and RNA modification detection via fast and accurate signal alignment"

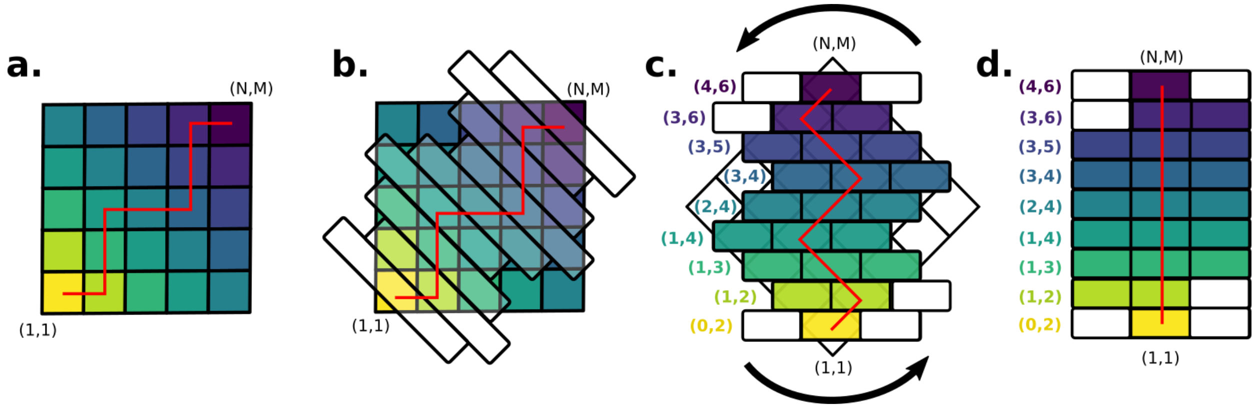


**Supplemental Figure 1.** Illustration of basecaller-guided DTW. The red line in each panel represents a basecalled alignment projected into signal space, which guides the band placement, determining which cells get computed. (a) A standard NxM DTW matrix, where N=M=5. Cells are colored by their Manhattan distance from (1,1), which corresponds to the band which they will be contained in. (b) The same DTW matrix overlaid with bands centered on the basecalled alignment (band width W=3). (c) The DTW band matrix with each row offset by its location in the NxM matrix, which is shaded in the background and rotated 45o. White cells indicate out-of-bounds coordinates. Band start coordinates are indicated by the colored numbers to the left. (d) The DTW band matrix, represented as a standard two-dimensional array. Note that the start coordinates are required to reconstruct the original matrix structure.

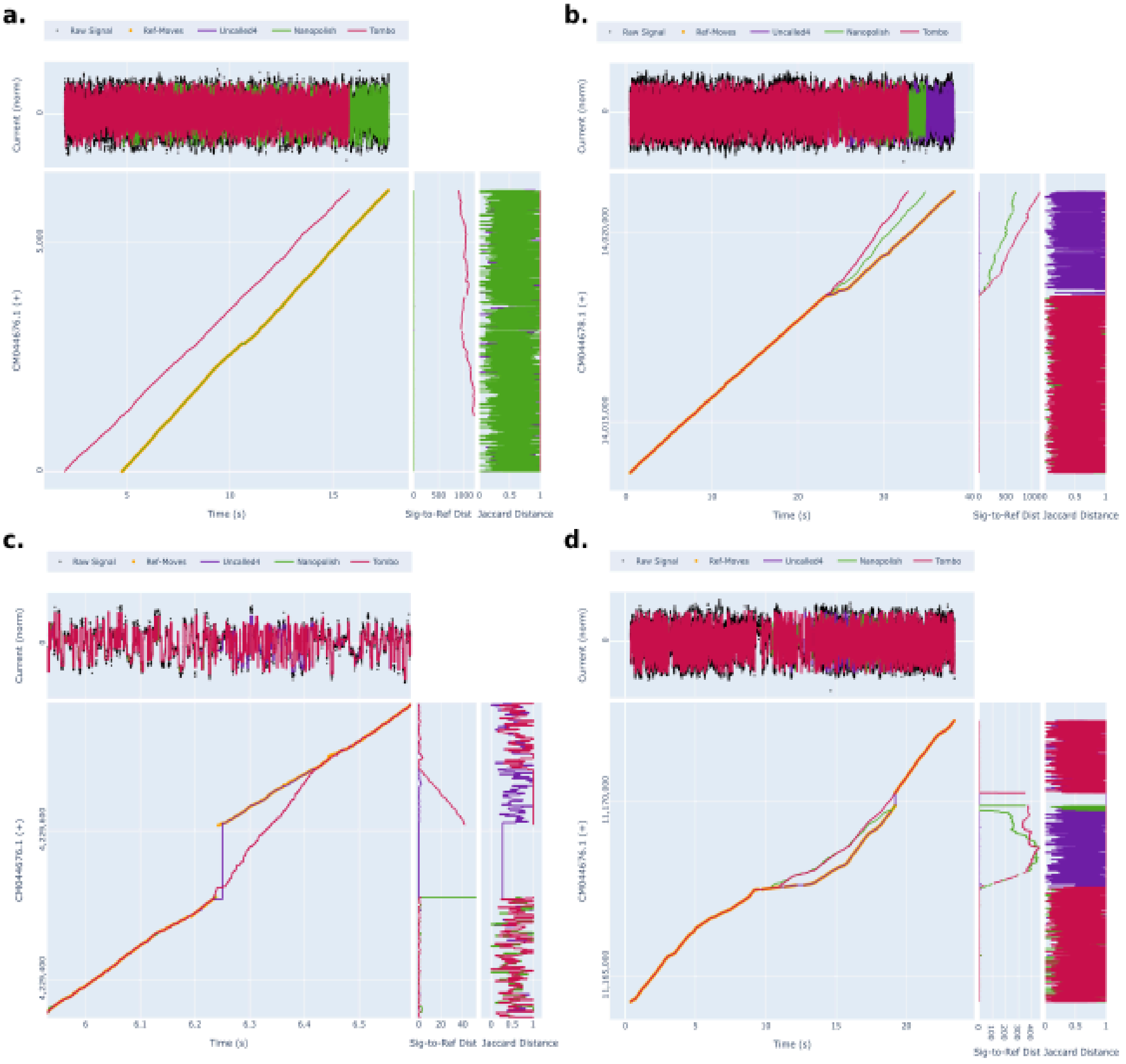


**Supplemental Figure 2.** Large-scale alignment errors in *D. melanogaster* DNA sequenced with r9.4.1. **(a)** An intergenic repeat causing Tombo to be shifted by several hundred bases. **(b)** A read which appears to change speed at the end, causing all three signal aligners to output different alignment endpoints, with Uncalled4 remaining close to the basecaller *ref-moves.* **(c)** A large deletion which Tombo aligns over, while Nanopolish and Uncalled4 properly skip the region. **(d)** A read with inconsistent speed, causing internal disruptions in Tombo and Nanopolish alignment.

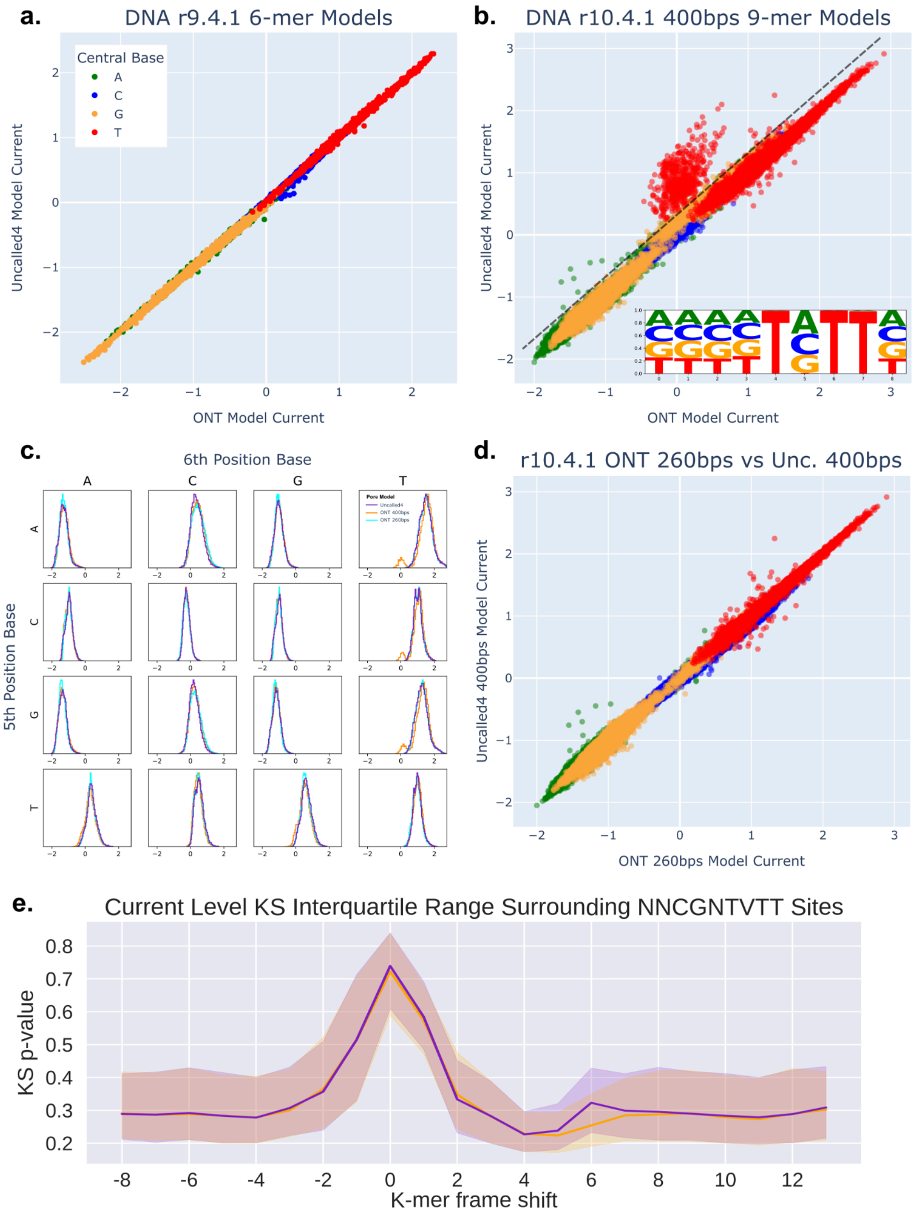


**Supplemental Figure 3.** DNA model training results. **(a)** Current levels from Uncalled4 and ONT r9.4.1 6-mer DNA models. **(b)** Current levels from Uncalled4 and ONT r10.4.1 400bps 9-mer DNA models. Inset displays sequence logo for k-mers with more than 0.5 normalized units of difference between the models (indicated on main plot by dashed line). **(c)** Current distributions for k-mers with each base fixed at the 6th and 5-th positions for r10.4.1 models, including both 400bps and 260bps ONT models. Most distributions are unimodal, except for ONT 400bps which has outliers caused by “TVTT” k-mers. **(d)** Comparison between Uncalled4’s r10.4.1 400bps model and ONT’s 260bps model, which lacks the outliers seen in ONT’s 400bps model. **(e)** KS statistics surrounding 5mCpG sites in TVTT context using Uncalled4 (purple) and ONT (orange) r10.4.1 400bps models.

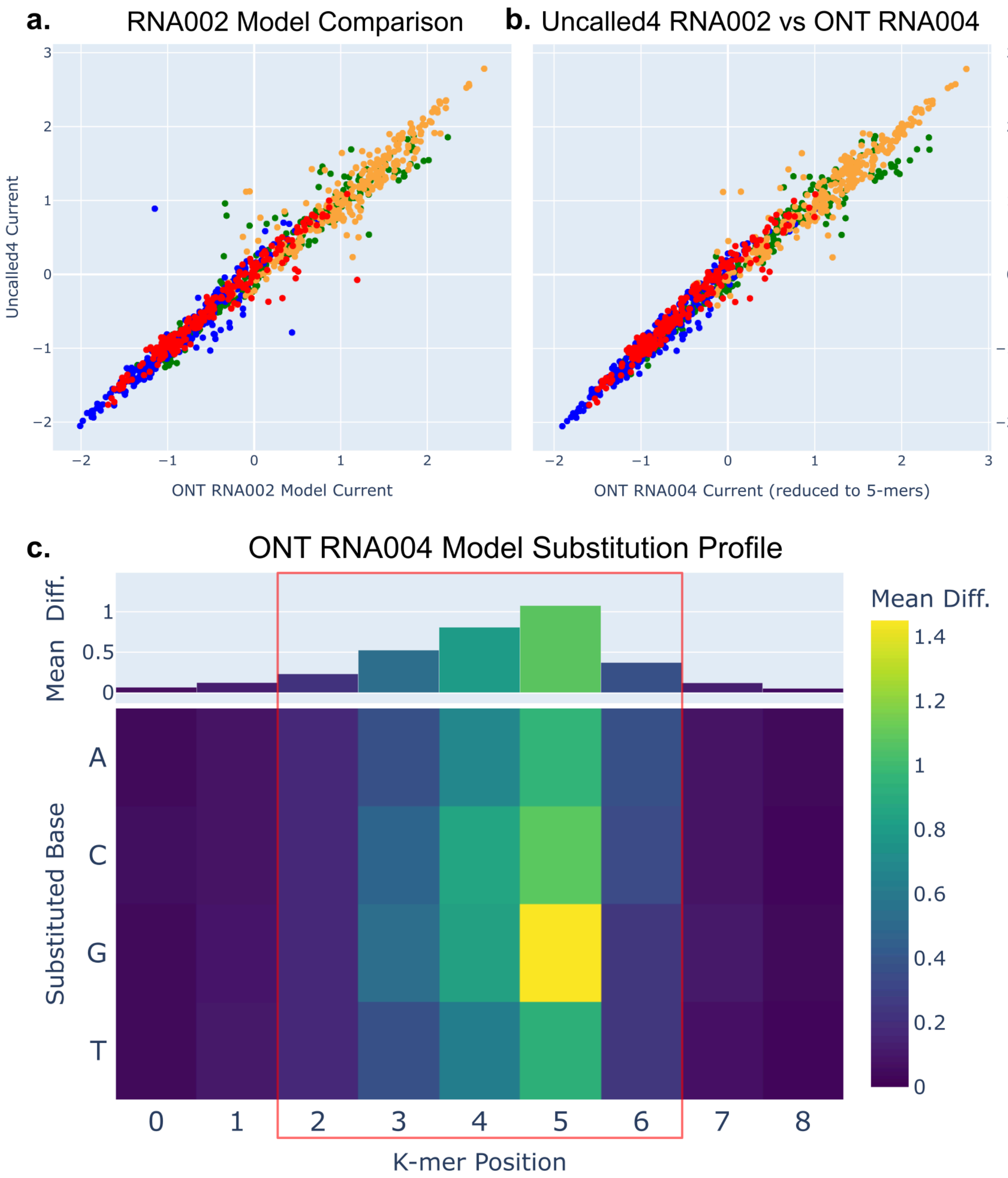


**Supplemental Figure 4.** RNA model training results. **(a)** Uncalled4 and ONT r9.4.1 RNA002 model comparison. **(b)** Comparison between the five central bases of ONT's RNA004 model and Uncalled4's RNA002 model. **(c)** Substitution profile of ONT's RNA004 model, with the five central bases highlighted to note the similarity with the RNA002 model (**Fig. 1a).**

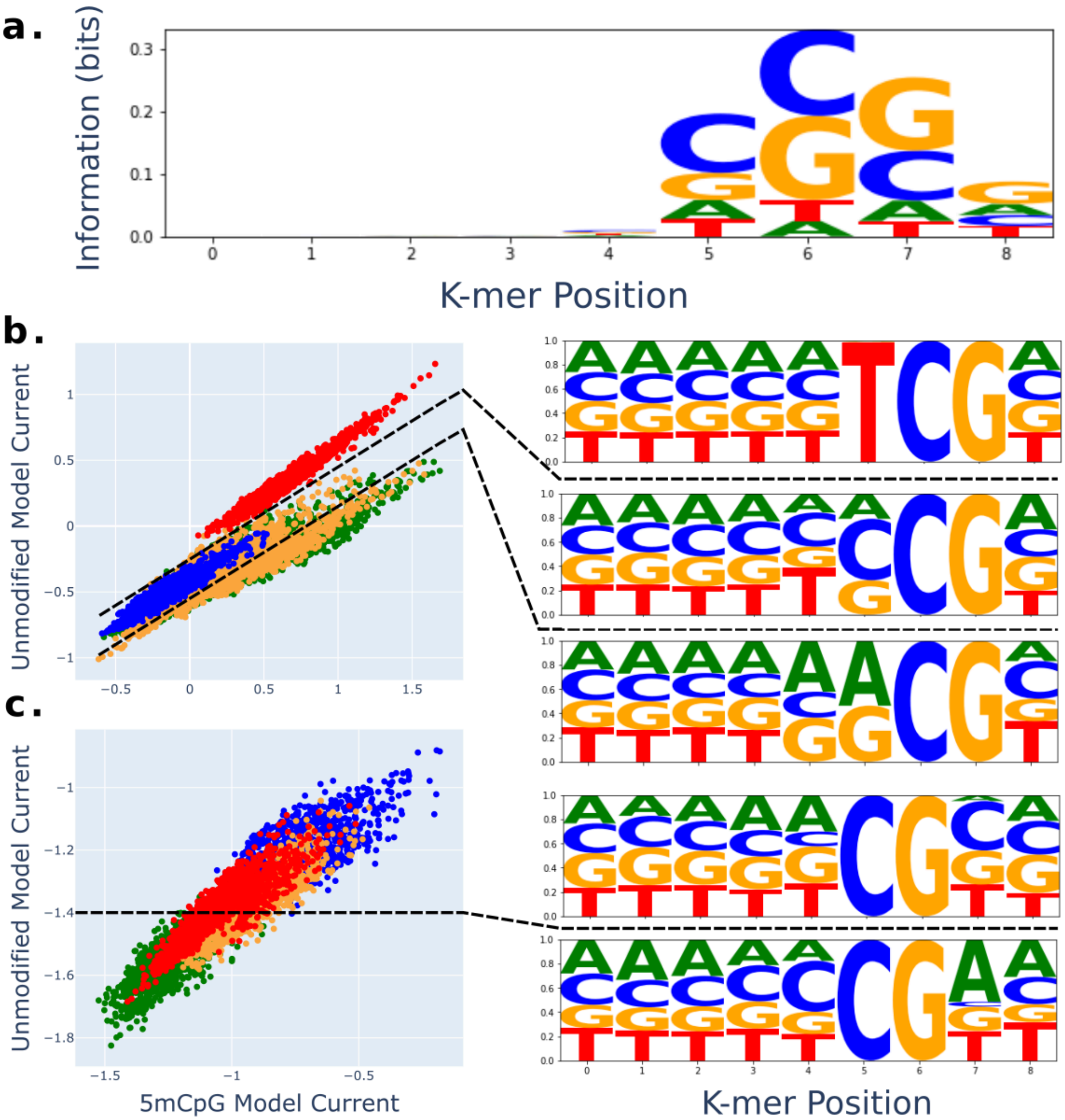


**Supplemental Figure 5.** 5mCpG model analysis. For comparison, the CpG model currents were linearly scaled using linear regression between non-CpG k-mers in Uncalled4's unmodified and 5mCpG models. **(a)** Per-base information content of k-mers with higher than median deviation between unmodified and 5mCpG models, showing CpG in any of the last four positions provides the most information. **(b)** Unmodified and 5mCpG currents for k-mers with CG in 6th and 7th positions, colored by the identity of the 5th position, alongside sequence logos for three subsets of k-mers divided by dashed lines. **(c)** Similar to (b) but for k-mer with CG in the 5th and 6th position, colored by 6th position identity and sequence logos for two subsets.

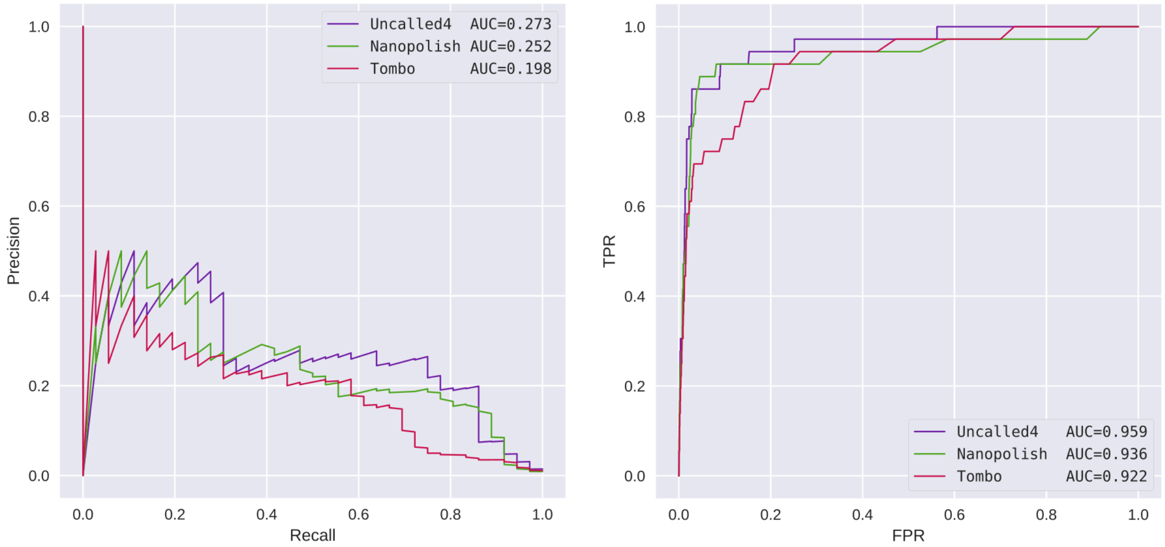


**Supplemental Figure 6.** Precision recall (left) and ROC (right) curves for detecting 36 modifications in *E. coli* ribosomal RNA, using KS statistics comparing current levels from native in *in vitro* transcribed RNA.

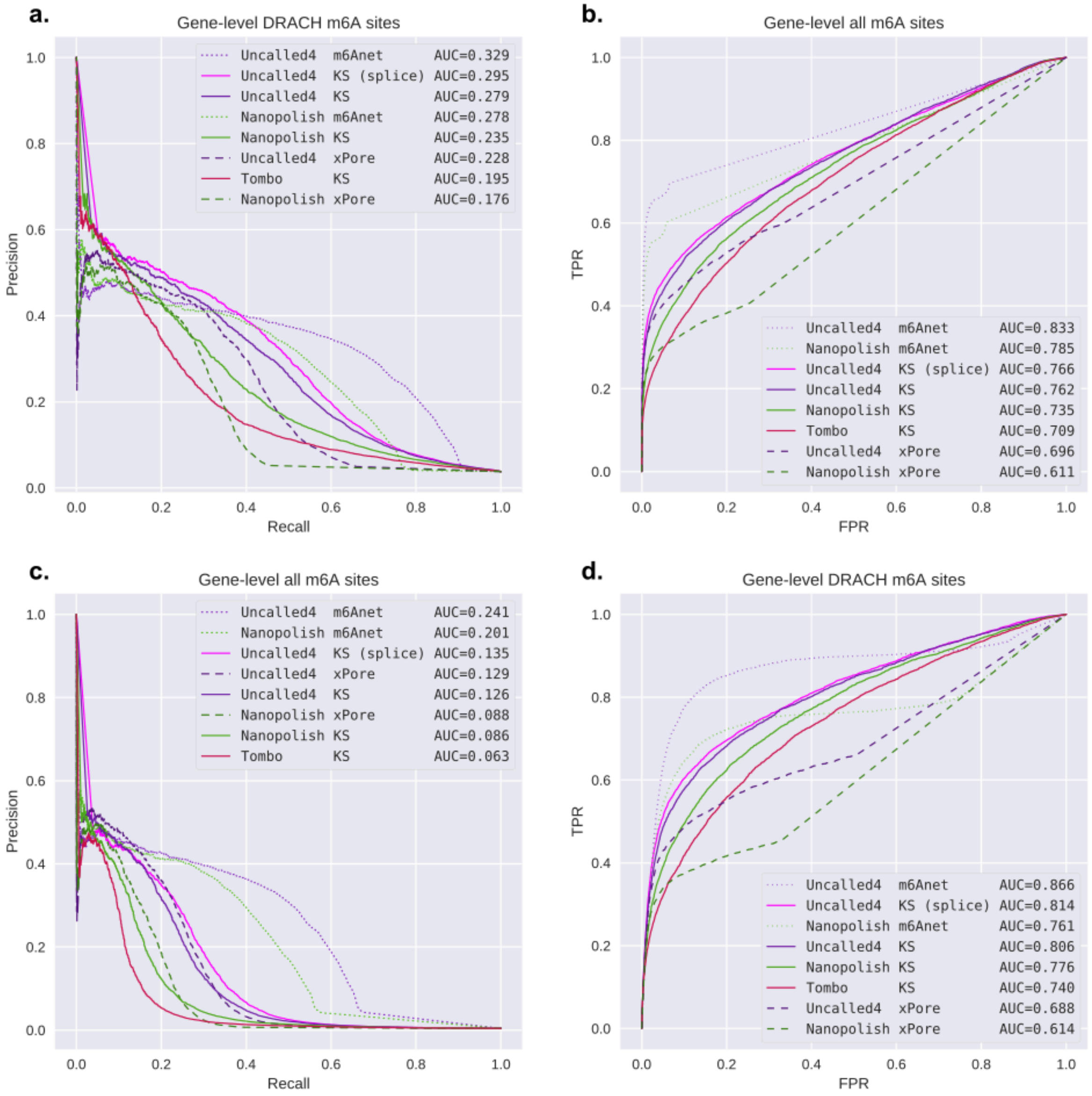


**Supplemental Figure 7.** Precision recall and ROC curves for transcript-level comparative m6A detection in HEK293t in all contexts (**a-b**) and limited to DRACH sites (**c-d**).

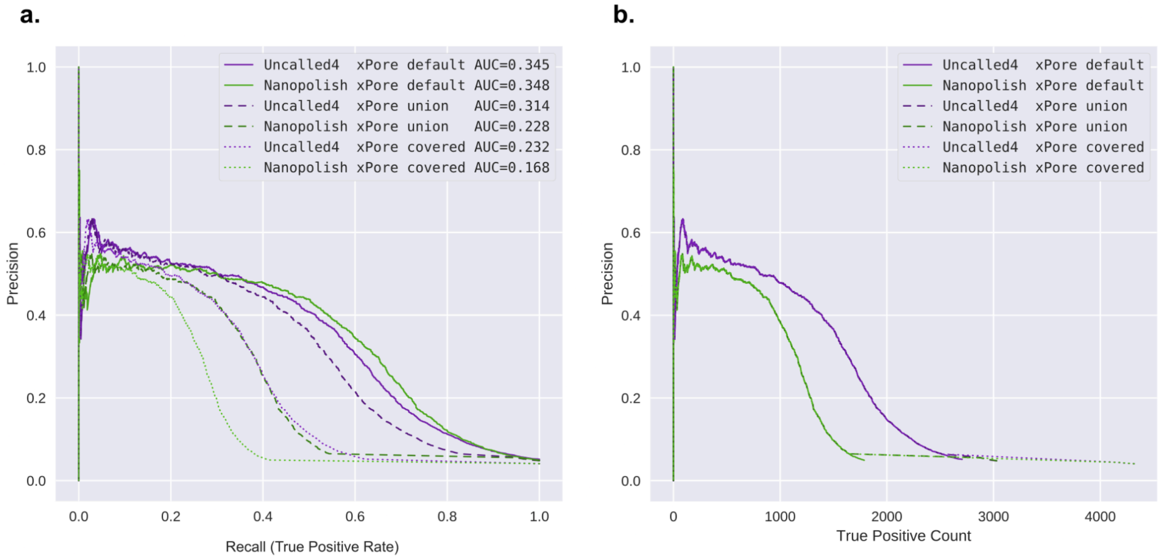


**Supplemental Figure 8. (a)** Precision recall curves for transcript-level m6A detection in HEK293t with xPore results, using three different methods for handling sites not output by xPore: "default" is naive comparison where missing values are ignored and the output sites and directly input to the precision/recall curve; "union" takes the union of all sites output by each aligner, and assigns a score of "0" for sites not output by either aligner, yielding a lower apparent recall for Nanopolish due to more missing data; "covered" includes all sites covered by minimap2 alignments at least 20x coverage, again filling in a score of "0" for sites not output by xPore. **(b)** Similar to a precision recall curve, but with the absolute true positive count rather than the rate (recall) on the x-axis, causing all three missing data filling strategies to output identical curves.

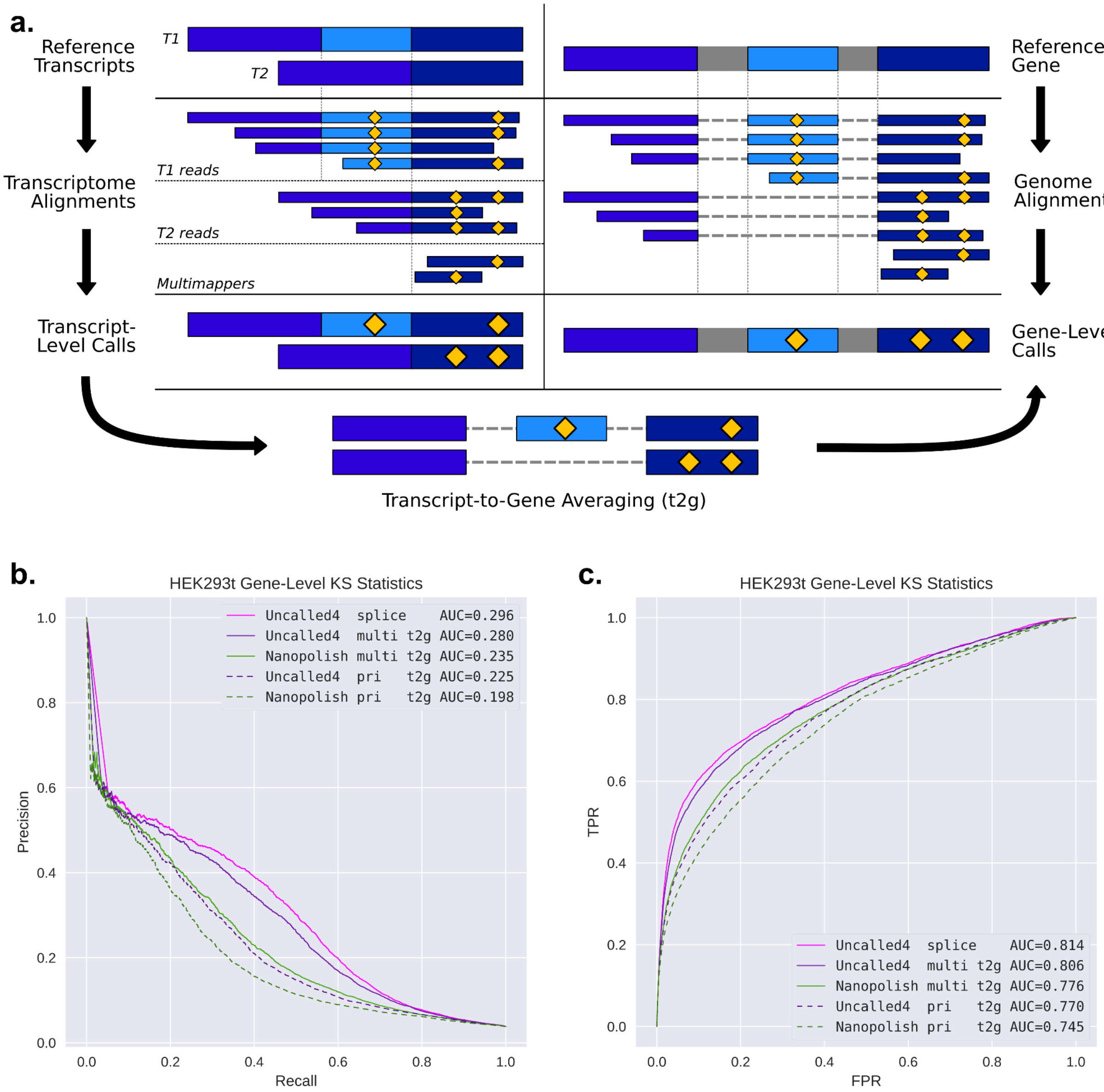


**Supplemental Figure 9. (a)** Illustration of transcript-level modification calling, genome-level calling, and translation of transcript-level calls to the gene-level (t2g). **(b)** Precision-recall and **(c)** ROC curves of Uncalled4 and Nanopolish gene-level calls using KS statistics. “Splice” indicates Uncalled4 spliced genome alignment. “Multi t2g” indicates transcript-to-gene averaging using all multi-mapping reads, while “pri t2g” indicates the same but only using primary alignments.

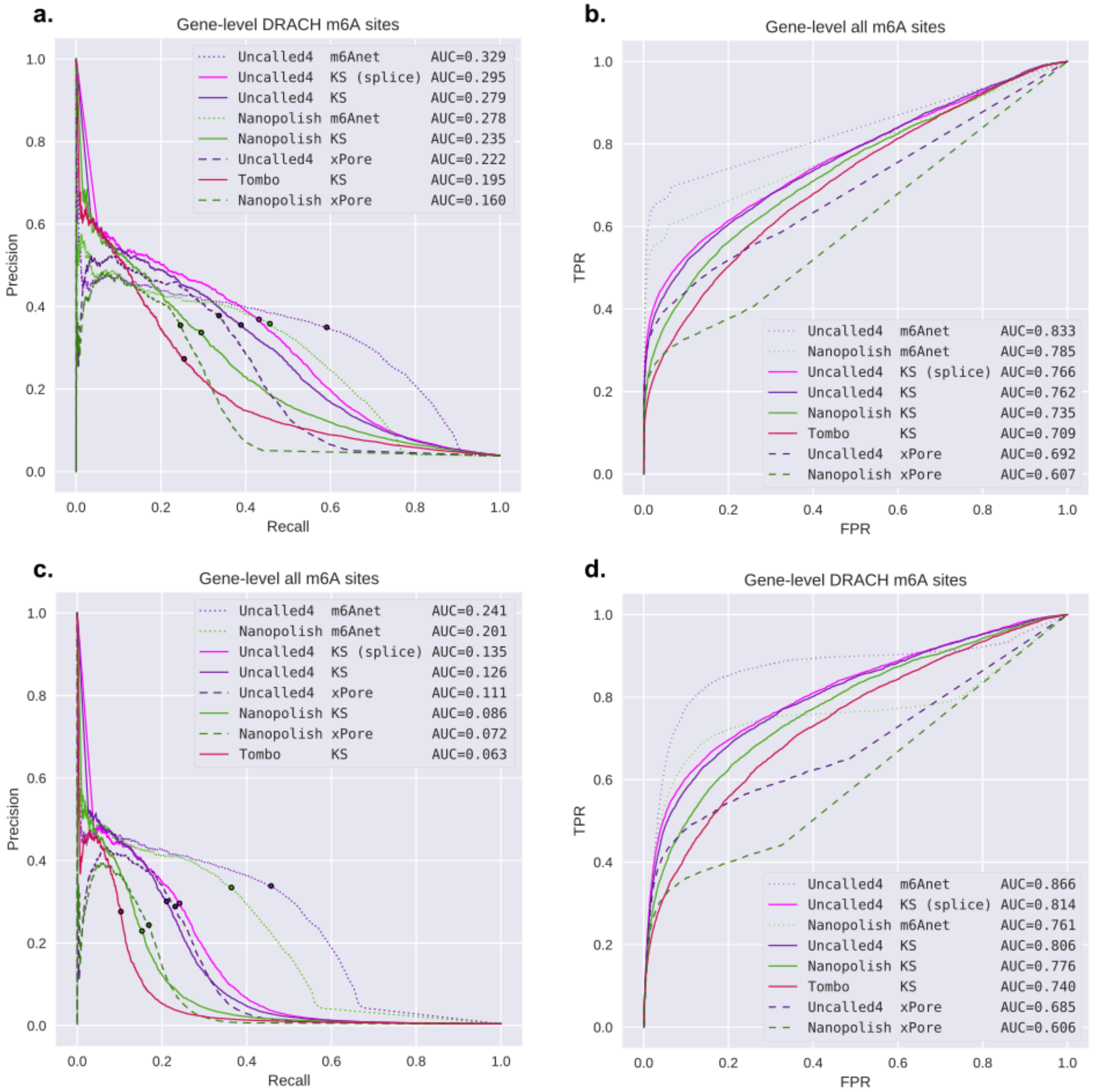


**Supplemental Figure 10** Precision recall and ROC curves for transcript-level comparative m6A detection in HEK293t in all contexts (**a-b**) and limited to DRACH sites (**c-d**). “splice” indicates Uncalled4 spliced genome alignment. All other methods used transcriptome alignments with all multi-mappers included, averaged to the gene-level.

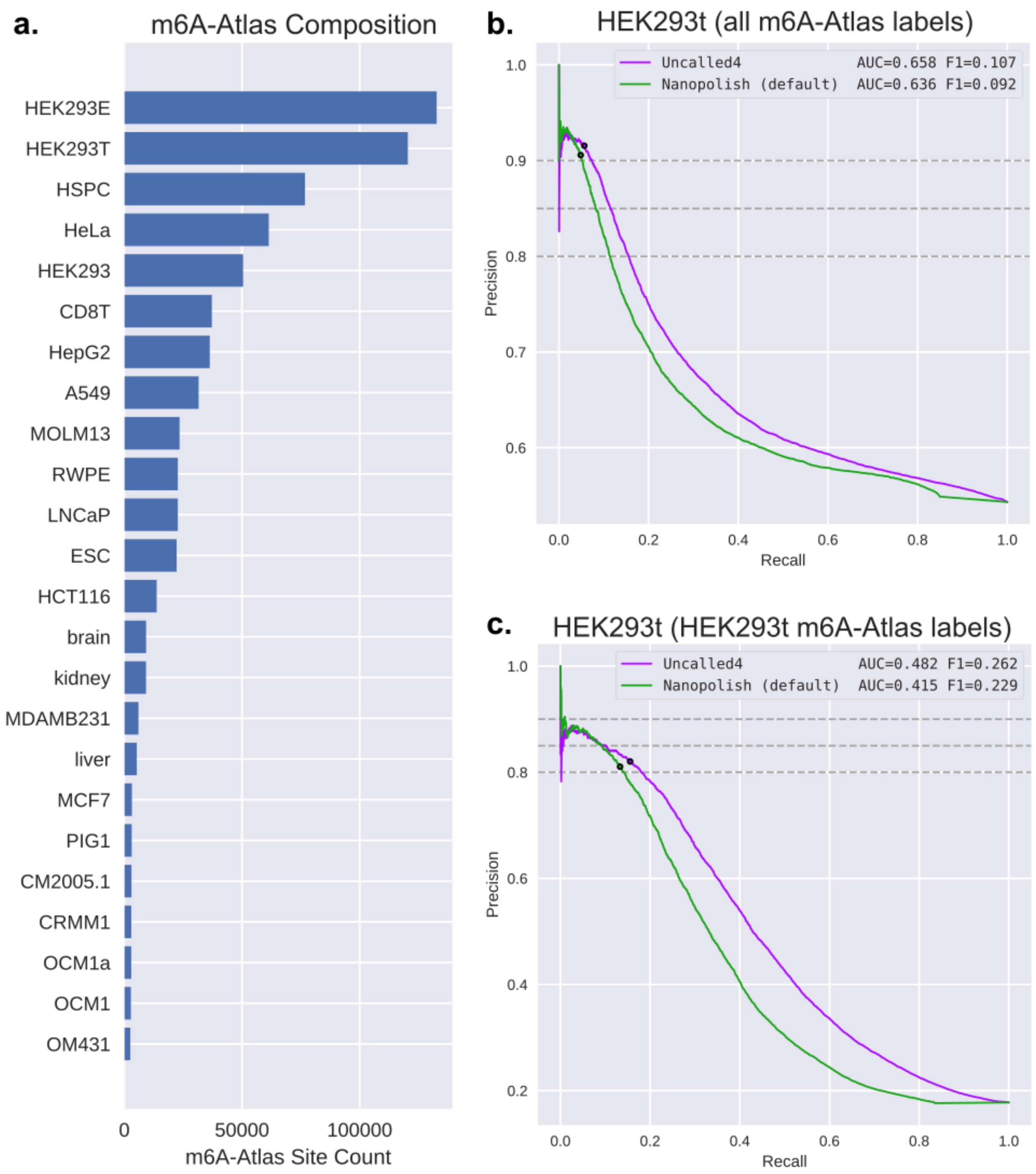


**Supplemental Figure 11** (**a**) Number of sites from each cell line or tissue in the m6A-Atlas version 2. Many sites occur in multiple samples, meaning they are counted twice here. (**b**) Precision-recall curve of transcript-level m6Anet calls on a single HEK293t replicate using all m6A-Atlas sites as the set of truely modified sites. (**c**) Precision-recall curve of transcript-level m6Anet calls on a single HEK293t replicate only using sites labeled with HEK293t in the m6A-Atlas.

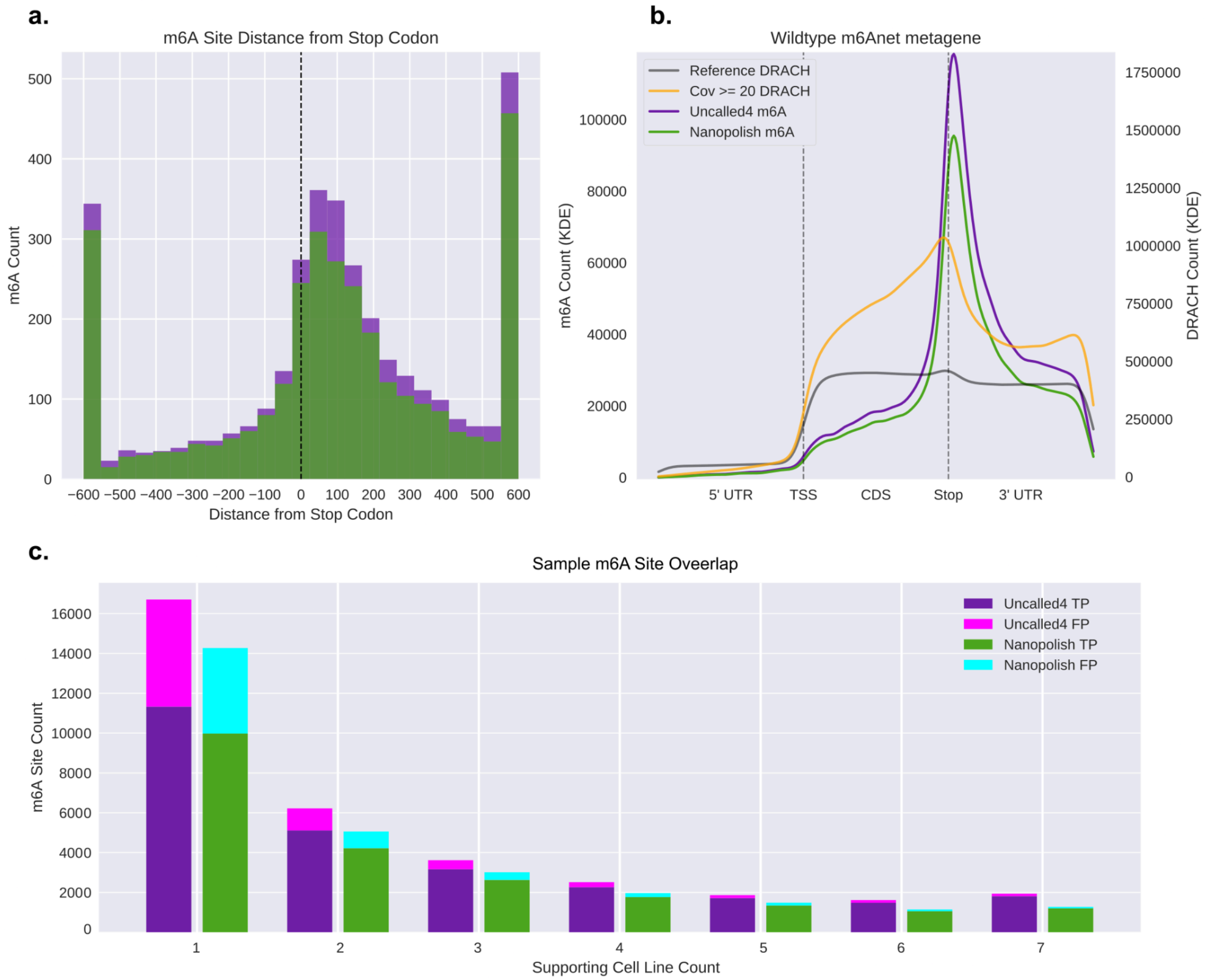


**Supplemental Figure 12. (a)** Distance from annotated stop codon for transcript-level m6A sites found by Uncalled4 (purple) and Nanopolish (green) with m6Anet at matched 85% precision. **(b)** Metagene plot of the same m6A sites, with the distribution of reference DRACH sites (gray) and DRACH sites covered by the nanopore reads (orange). **(c)** Gene-level m6A counts by the number of cell lines they were found in, divided into putative “true positives” (TP, in the m6A-Atlas) and putative “false positives” (FP, missing from the m6A-Atlas).
